# Differential binding of salicylic acid, phenolic acid derivatives and co-factors determines the roles of Arabidopsis NPR1 to NPR4 in plant immunity

**DOI:** 10.1101/2022.02.18.481035

**Authors:** Evelyn Konopka, Mathias Saur, Artur J.P. Pfitzner, Ursula M. Pfitzner

## Abstract

**Summary:**

- Genetic studies have demonstrated that NPR1 is the key positive regulator of salicylic acid (SA)-induced *PR-1* gene induction and systemic acquired resistance (SAR). In Arabidopsis, family members NPR1 to NPR4 share domain architecture.
- Yeast hybrid assays were used to explore biochemical capabilities of NPR1 to NPR4.
- All NPR1 to NPR4 are responsive to SA. SA perception proceeds via the conserved arginine embedded in a C-terminal LENRV-like motif. Clade 2 proteins NPR3 and NPR4 perceive SA directly, while clade 1 members NPR1 and NPR2 require interaction with partner proteins NIMIN1/NIMIN2 and TGA factors, respectively, to enable SA sensing. Intriguingly, NPR3 is considerably more sensitive to the synthetic analog 3,5-dichloroanthranilic acid than to SA, and all NPR1 to NPR4 are able to sense the microbial metabolite 6-methyl SA.
- We suggest that the plant´s ability to track SA and phenolic acid derivatives through NPR proteins has evolved to support diverse defense signaling outputs that are activated in parallel by agonists which may be of microbial or plant origin. In this line, NPR1-NIMIN2/NIMIN1 complex is the prime receptor for SA synthesized by plants in response to microbial attack, while NPR3 induces defense different from SAR primarily via unrecognized signal molecules.

## Introduction

Systemic acquired resistance (SAR) is a broad-spectrum and long-lasting defense response induced by effector-triggered immunity that prevents spread of pathogens within plants and protects them from secondary infection (Ross, 1961; Fu & Dong, 2013). SAR is activated by the signal molecule salicylic acid (SA; Malamy *et al*., 1990; Métraux *et al*., 1990; Vlot *et al*., 2009) leading to accumulation of PATHOGENESIS-RELATED (PR) proteins with antimicrobial activities (Ward *et al*., 1991; van Loon *et al*., 2006). Induction of *PR-1* genes serves as marker for SAR. Phenotypically, SAR is inconspicuous and has been associated with cell survival (Zavaliev *et al*., 2020).

The central regulator of SAR is *NON-EXPRESSOR OF PR GENES1* (*NPR1*), a.k.a. *NON-INDUCIBLE IMMUNITY1* (*NIM1*) and *SALICYLIC ACID INSENSITIVE1* (*SAI1*; Cao *et al*., 1994; Delaney *et al*., 1995; Cao *et al*., 1997; Ryals *et al*., 1997; Shah *et al*., 1997). The gene was identified through different genetic screens from Arabidopsis mutants. The Arabidopsis *NPR1* gene codes for a 66 kDa protein with known protein-protein interaction motifs. Accordingly, yeast two-hybrid screens identified interaction partners for NPR1.

TGACG-BINDING FACTORS (TGAs) link NPR1 with SA-responsive *as-1*-like *cis*-acting elements present in promoter regions of Arabidopsis (At) and tobacco (Nt) *PR-1* genes (Lebel *et al*., 1998; Strompen *et al*., 1998; Zhang *et al*., 1999; Després *et al*., 2000; Niggeweg *et al*., 2000; Zhou *et al*., 2000). Thus, NPR1 functions as co-activator of SA-induced *PR-1* gene transcription (Rochon *et al*., 2006). Another group of NPR1 binding partners are NIM1-INTERACTING (NIMIN, N) proteins (Weigel *et al*., 2001). In Arabidopsis, there are four *NIMIN* genes, *N1, N1b, N2* and *N3*. Of these, *NIMIN1* and *NIMIN2* are themselves activated by SA, and overexpression of *NIMIN1* represses SA induction of *PR* genes (Weigel *et al*., 2005; Hermann *et al*., 2013). Both SA-induced NIMIN1 and NIMIN2 interact strongly with a region in the conserved C-termini of Arabidopsis and tobacco NPR1, termed N1/N2-binding domain (N1/N2BD; Maier *et al*., 2011).

*NPR1* is eponym for a small gene family comprising six members in Arabidopsis. Three paralogs, NPR2, NPR3 and NPR4, exhibit high homology to NPR1, in particular in the proteins´ C-terminal thirds. Paralogs NPR3 and NPR4, like NPR1, have also been associated with pathogen defense reactions (Liu *et al*., 2005; Zhang *et al*., 2006). However, only NPR1 has been identified as positive regulator of SA-induced *PR-1* gene expression and SAR through different genetic screens. In one model, NPR3 and NPR4 control levels of NPR1 in an SA-dependent fashion via proteasomal degradation thus promoting SAR at intermediate SA concentrations (Fu *et al*., 2012). Other work suggested that NPR3 and NPR4 negatively regulate *PR* gene expression and pathogen resistance (Zhang *et al*., 2006; Shi *et al*., 2013). Recently, it has been postulated that NPR3 and NPR4 act redundantly as transcriptional co-repressors of SA-responsive genes independently from NPR1, and that NPR3/NPR4-mediated defense gene repression is relieved by SA (Ding *et al*., 2018).

Since its discovery in 1997, different models have been proposed for transduction of the SA signal through NPR1. The most robust model was initially issued for tobacco NPR1 based on various yeast hybrid (YH) assays (Maier *et al*., 2011; Neeley *et al*., 2019). Two conserved C-terminal domains of NtNPR1, the LENRV motif and the N1/N2BD region, separated from each other, were shown to physically associate in yeast driven by SA. Physical association of the two domains results in re-organization of the C-terminus yielding enhanced transcription activation through NtNPR1. The arginine in the LENRV motif is mandatory for SA sensing. This model is supported by biochemical and genetic evidence obtained for Arabidopsis NPR1 (Ryals *et al*., 1997; Rochon *et al*., 2006; Canet *et al*., 2010) and Arabidopsis NPR4 (Ding *et al*., 2018). Furthermore, the key features, namely re-modeling of the C-terminus by SA-mediated association of two distinct domains, the LENRV-like and the N1/N2BD regions, have recently been confirmed for SA-bound Arabidopsis NPR4 by X-ray crystallography (Wang *et al*., 2020).

To further explore SA signaling capacities of Arabidopsis NPR proteins sharing the NPR1 domain structure, we compared biochemical capabilities of NPR1 to NPR4. We used various YH assays developed by us, in particular N1/N2-NPR interaction and the split LENRV-N1/N2BD Y2H assay. We show that all Arabidopsis NPR1 to NPR4 sense the SA signal molecule via the conserved arginine residue in a LENRV-like motif. Yet, NPR1 to NPR4 exhibit different protein-protein interaction preferences and different sensitivities to SA and structurally related compounds. Our findings support the notion of differential activation of the SA sensors in the Arabidopsis NPR family producing distinct signaling outputs.

## Materials and Methods

### DNA constructs

For protein-protein interaction assays in yeast, cDNA sequences encoding truncated or full-length proteins were fused in-frame to the sequence for Gal4 DNA-binding domain (GBD) in pGBT9 and the sequence for Gal4 transcription activating domain (GAD) in pGAD424. Arabidopsis and tobacco *NPR1* clones as well as *NIMIN, TGA2* and *TGA6* constructs were described previously (Weigel *et al*., 2001; Neeley *et al*., 2019). Generally, clones were generated by PCR amplification using primers listed in Tables S1 to S3. Details for construction of clones encoding TGA factors and full-length, partial and mutant NPR proteins are described in Methods S1. Subregions and mutants of NPR proteins analyzed in YH assays are depicted in Fig. S1. All clones generated by PCR amplification were verified by DNA sequence analysis.

### Yeast hybrid analyses

Yeast hybrid assays (Y1H, Y2H, Y3H) were conducted as reported earlier (Weigel *et al*., 2001; Maier *et al*., 2011; Neeley *et al*., 2019). Yeast cells were grown in absence or presence of different chemicals as indicated. Compounds insoluble in water were solved in dimethyl sulfoxide (DMSO). The final DMSO concentration in yeast growth medium was from 0.1% to 0.5%. *LacZ* reporter gene activities were tested in duplicate with at least three independent colonies. Experiments were repeated at least once with new yeast transformants.

Representative results collected in one experimental set are shown. Results are depicted as mean enzyme activities in Miller units, plus and minus standard deviation (SD).

### Chemical treatment of plants

Tobacco (*N. tabacum* cv. Samsun NN) and Arabidopsis Col-0 plants were grown in commercial potting soil under natural light conditions in a greenhouse. For chemical treatment, leaf disks were cut from young tobacco plants and floated for 4 days on water or solutions of 1mM SA, acetylsalicylic acid (Ac-SA), 5-fluorosalicylic acid (F-SA), 6-methylsalicylic acid (6-MeSA) or 2,5-dihydroxybenzoic acid (2,5-diOH BA) containing 0.3% DMSO. Concentration of 3,5-dichloroanthranilic acid (3,5-DCA) was 0.1mM in 0.3% DMSO. Protein extraction and GUS reporter enzyme assays were performed as described previously (Stos-Zweifel *et al*., 2018). GUS activities are given in units (1 unit = 1nMol 4-MU formed per hour per mg protein).

### Immunodetection of NPR1

Immunodetection was performed with a polyclonal antiserum raised against 6xHis–AtNPR1 as described by Stos-Zweifel *et al*. (2018). NPR1 accumulation in Arabidopsis Col-0 was analyzed 14 hr after spraying plants with H_2_O, 1mM SA or Bion containing 0.3mM BTH. Equal amounts of leaf tissue were extracted with GUS lysis buffer. Crude extracts were cleared twice by centrifugation, and equal extract volumes were loaded on a 10% SDS gel as indicated. Protein loading was checked by staining the nitrocellulose filter with Ponceau S (0.1% in 5% acetic acid). For detection of GBD–NPR1 fusion protein in yeast, extracts were prepared from transformed cells as reported by Weigel *et al*. (2001). An unspecific band reacting with the antiserum is marked for demonstration of equal gel loading.

### Bioinformatic tools

Phylogenetic analysis was performed on the Phylogeny.fr platform using the “One Click” software developed by Dereeper *et al*. (2010). Briefly, protein sequences were aligned with MUSCLE (v3.8.31). After alignment, ambiguous regions were removed with Gblocks (v0.91b). The phylogenetic tree was reconstructed using the maximum likelihood method implemented in the PhyML program (v3.1/3.0 aLRT). Graphical representation and edition of the phylogenetic tree were performed with TreeDyn (v198.3).

## Results

### Domain structure of Arabidopsis NPR1 to NPR4 and gene expression

Arabidopsis NPR family members are organized in three clades, clade 1 (NPR1 and NPR2), clade 2 (NPR3 and NPR4) and clade 3 (BOP1 and BOP2; Fig. 1a). Each clade contains two highly related members believed to have evolved after gene duplication from a common ancestral gene. NPR1 to NPR4 share domain structure including an N-terminal broad complex, tramtrack, and bric à brac/poxvirus and zinc finger (BTB/POZ) domain, a central ankyrin repeat domain and a similar C-terminal region with a LENRV-like motif and a conserved N1/N2BD (Fig. 1b). While Arabidopsis NPR1, like tobacco NPR1, harbors the signature LENRV, AtNPR2 contains the sequence YENRV, and AtNPR3 and AtNPR4 both contain the sequence LEKRV (Fig. S1a). Of note, AtNPR2 and AtNPR3 also feature the critical arginine previously associated with sensing and binding SA in NPR1 and NPR4 (Maier *et al*., 2011; Ding *et al*., 2018; Neeley *et al*., 2019; Wang *et al*., 2020).

**Fig. 1.**
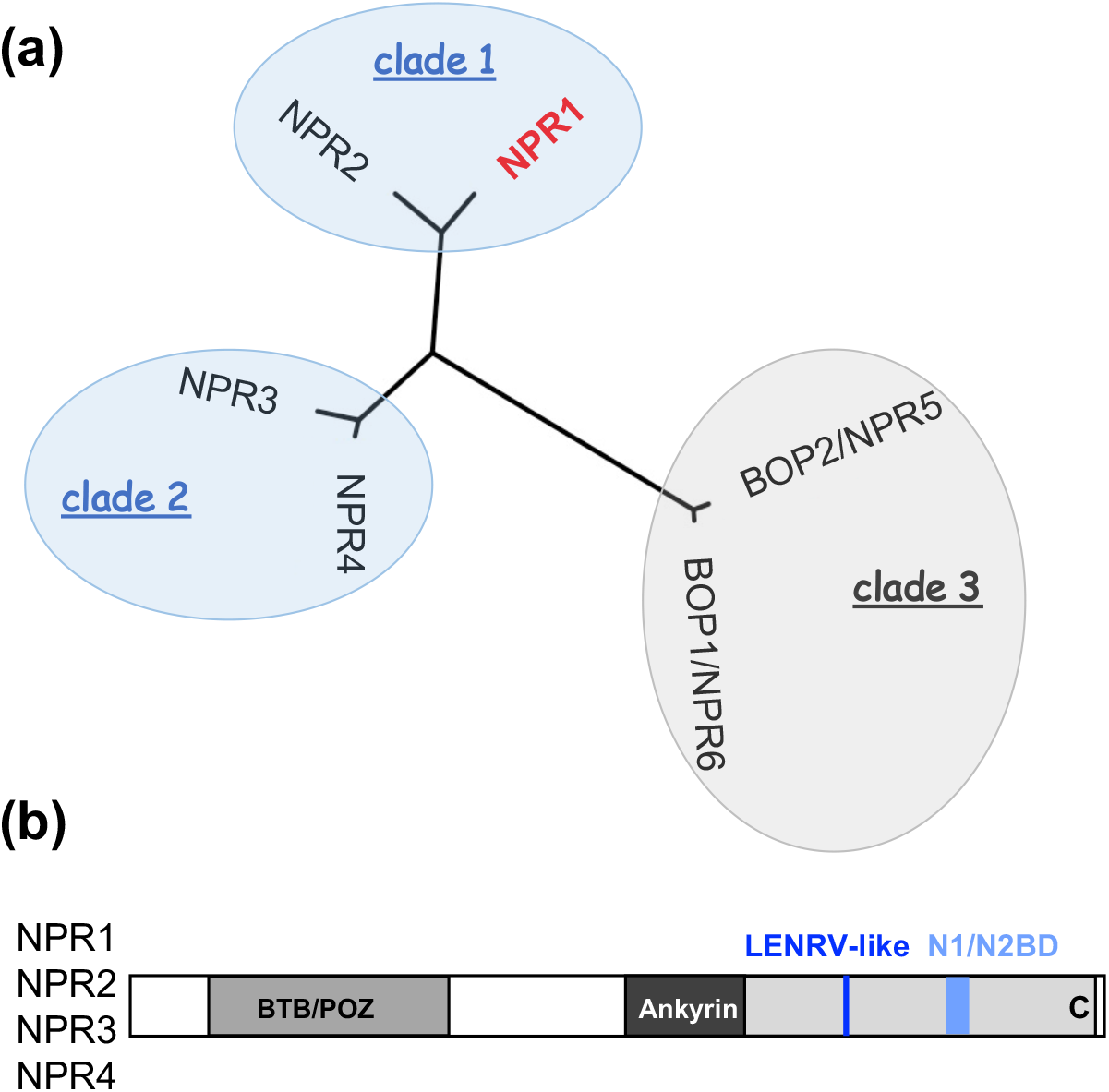
Relatedness of Arabidopsis NPR1 to NPR4. (a) Unrooted phylogenetic tree of Arabidopsis NPR family proteins. The tree was constructed with a software developed by Dereeper *et al*. (2010). The six NPR proteins fall in three clades with two highly similar members in each clade. (b) Domain structure of Arabidopsis NPR1 to NPR4. Conserved domains present in NPR1 to NPR4 are a BTB/POZ domain, an ankyrin repeat region and a C-terminal NPR1-like sequence (C, in light grey) with a LENRV-like motif and a NIMIN1/NIMIN2-binding domain (N1/N2BD) consensus sequence.

*NPR1, NPR3* and *NPR4* are all expressed to intermediate levels in Arabidopsis leaves while *NPR2* is expressed only rather low (Shi *et al*., 2013; Klepikova *et al*., 2016). Consistently, NPR1 protein accumulated in non-treated leaf tissue (Fig. S2). After exposure of plants to SA or its functional analog benzothiadiazole (BTH), protein accumulation was enhanced moderately. Collectively, these findings are compatible with the role of NPR1 as SA sensor protein.

### Transcription activity of Arabidopsis NPR1 to NPR4 in yeast

To evaluate biochemical capabilities of NPR1 to NPR4 in SA signaling, we cloned the cDNAs from Arabidopsis Col-0 and compared activities of the proteins using various YH assays which we developed previously for characterization of tobacco NPR1 and which led us to predict the mechanism of SA perception via the two conserved C-terminal domains LENRV and N1/N2BD (Maier *et al*., 2011; Neeley *et al*., 2019). Although NtNPR1 displays low level transcription activity in Y1H assays that is enhanced by SA, we were not able to measure substantial autoactivation potential with Arabidopsis NPR1 to NPR4 in presence or absence of SA (Fig. S3).

### Dimerization of Arabidopsis NPR1 to NPR4 in yeast

The model proposed by Fu *et al*. (2012) suggests nuclear accumulation of NPR1, and thus *PR-1* gene expression, only at intermediate levels of SA. At very low and high hormone concentration, NPR1 protein is degraded due to SA-controlled dimerization with NPR4 and NPR3, respectively, which function as CULLIN3 adaptor proteins. In addition, NPR3 and NPR4 homo and heterodimer formation was reported to be strengthened by SA (Fu *et al*., 2012; Palmer *et al*., 2019). The functional relevance of these latter observations remained elusive. We were not able to demonstrate NPR3 and NPR4 homo or heterodimerization nor NPR3/NPR4-NPR1 interaction in quantitative Y2H assays neither in absence nor in presence of SA (Fig. S4a-c). Immunodetection verified that GBD–NPR1 fusion protein accumulated in yeast cells co-transformed with pGAD424/NPR3 and pGAD424/NPR4 (Fig. S4d). Failure of NPR3/NPR4-NPR1 binding in Y2H has also been reported by Castelló *et al*. (2018) and Ding *et al*. (2018). Alone GBD–NPR2 fusion protein produced some reporter activity in dimerization experiments (Fig. S4b). Activities were, however, barely above background and did not bear comparison with our interaction standard GBD–NIMIN1 + GAD–NPR1. Furthermore, positive interactions of NPR2 described below were considerably stronger.

### Interaction of Arabidopsis NPR1 to NPR4 with TGA transcription factors in yeast

NPR1 constitutively binds basic leucine zipper (bZIP) TGA factors from clade II (TGA2,5,6) and clade III (TGA3,7). Accordingly, we found clear interaction of NPR1 with TGA2, TGA3 and TGA7. Typically, interaction between TGA2 and NPR1 was slightly stronger in presence of SA (Fig 2a; < 2-fold increase; Maier *et al*., 2011). Binding of clade II and clade III TGA factors occurred in the N-terminal two thirds of NPR1 encompassing the ankyrin repeat region (Fig. S5). We were not able to detect TGA factor binding to the C-terminal third of NPR1 (aa 387-593; data not shown). This is in contrast to tobacco NPR1 which uniquely binds transcription factor NtTGA7 near the N1/N2BD (Stos-Zweifel *et al*., 2018). Likewise, NPR3 and NPR4 interacted with TGAs of clades II and III in an overall less SA-affected fashion (Fig. 2c; < 2.5-fold change). The profiles of TGA factor binding were different for NPR1, NPR3 and NPR4. While NPR1 interacted about equally with all TGA2, TGA3 and TGA7, NPR3 interaction was strongest with TGA3, and NPR4 bound best to TGA2.

**Fig. 2.**
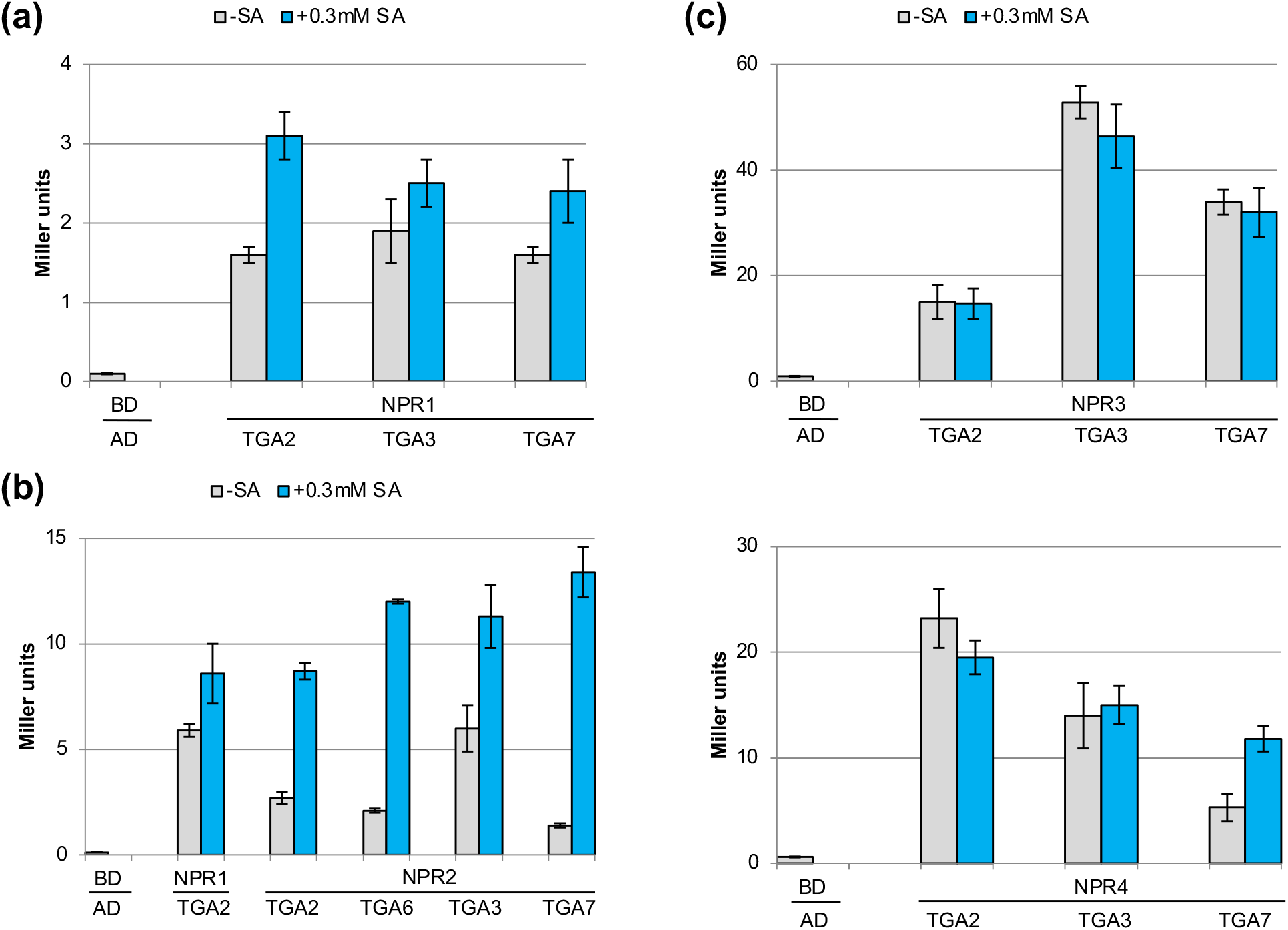
Interaction of Arabidopsis NPR1 to NPR4 with Arabidopsis TGA transcription factors in yeast. *NPR* genes were expressed in fusion with the sequence for Gal4 DNA-binding domain (BD), and *TGA* genes were expressed in fusion with the sequence for Gal4 transactivation domain (AD). Quantitative Y2H assays were conducted in absence and presence of salicylic acid. (a) Interaction of NPR1 with TGA2, TGA3 and TGA7. (b) Interaction of NPR2 with TGA2, TGA6, TGA3 and TGA7. (c) Interaction of NPR3 and NPR4 with TGA2, TGA3 and TGA7.

Similarly, NPR2 bound TGA factors of clades II and III constitutively. Quite surprisingly, however, NPR2-TGA interactions were generally enhanced in presence of SA. NPR2-TGA7 binding was most affected with an >3-fold increase by SA (Fig. 2b).

### Interaction of Arabidopsis NPR1 to NPR4 with NIMIN proteins in yeast

We also compared interaction of NPR1 to NPR4 with NIMIN proteins. We could not demonstrate binding of NIMIN1, NIMIN2 or NIMIN3 to NPR2, NPR3 or NPR4 in Y2H, neither in absence nor in presence of SA (data not shown). Our findings are in contrast to an earlier report (Altmann *et al*., 2020) and to strong interaction of NIMIN1 to NIMIN3 with NPR1 (Weigel *et al*., 2001). Consequently, NPR3 and NPR4, unlike NPR1, were unable to form ternary complexes with NIMIN2 and TGA transcription factors (Fig. S6). In summary, NPR1 to NPR4, although sharing domain structure, display differential protein-protein interaction potential indicating unique roles for N1/N2-NPR1 and TGA7-NPR2 complexes.

### Salicylic acid drives physical association of LEKRV and N1/N2BD regions of NPR3 and NPR4

Of note, failure of interaction between NIMIN1/NIMIN2 and NPR2 to NPR4 was against any expectation, as NPR2 to NPR4 contain C-terminal regions with high sequence conservation to the N1/N2BD identified by us in Arabidopsis and tobacco NPR1 (Fig. S1a). Recently, the sequences comprising the N1/N2BDs of tobacco NPR1 and Arabidopsis NPR4 have been implicated in yet another activity, i.e., association of LENRV-like and N1/N2BD regions via the SA signal molecule (Neeley *et al*., 2019; Wang *et al*., 2020). In fact, six residues within the NPR4 N1/N2BD as defined by us participate in forming the SA-binding pocket, and altogether ten out of fourteen residues identified in both the LENRV-like and N1/N2BD regions of NPR4, that are involved in binding SA, are shared by NPR1, NPR2 and NPR3 (Fig. S7; Wang *et al*., 2020). We therefore tested interaction potential of separated LENRV-like and N1/N2BD regions of Arabidopsis NPRs.

Unexpectedly, NPR1 was unable to associate its LENRV and N1/N2BD parts in presence of SA (Fig. S8a; for dimensions of LENRV-like and N1/N2BD regions of different NPR proteins used in Y2H, see Fig. S1b). Likewise, NPR2 did not show SA-induced activity with YENRV and N1/N2BD parts (Fig. S8b). On the contrary, C-terminal subregions of NPR3 and NPR4, respectively, displayed interaction when SA was added to yeast growth medium, albeit at a lower level as compared to tobacco NPR1 C-terminal parts (Fig. 3a). Association of NPR3 LEKRV and N1/N2BD parts did not occur in presence of the non-functional SA analog 4-hydroxybenzoic acid (4-OH BA; Fig. 3b).

**Fig. 3.**
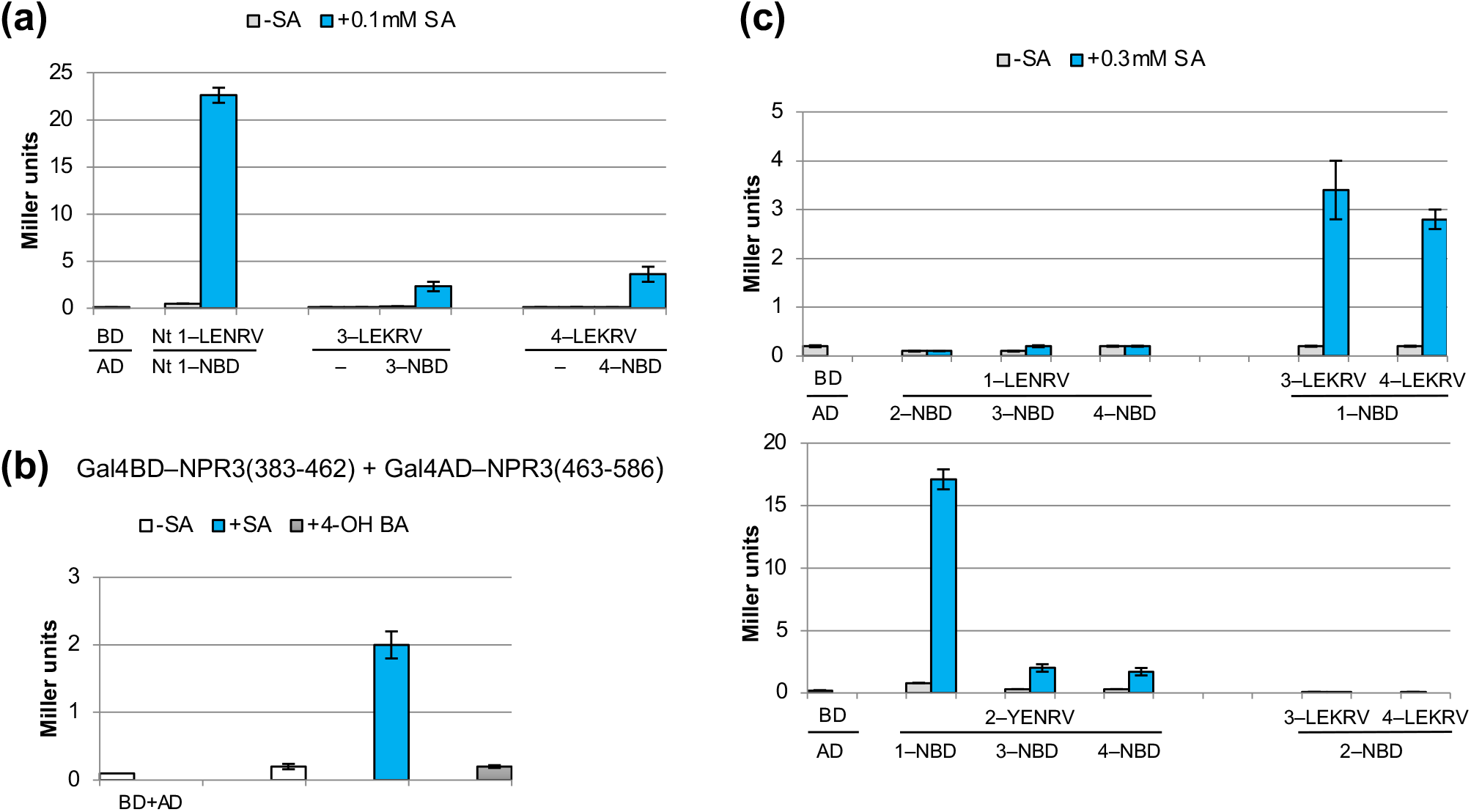
Interaction between separated LENRV-like and NIMIN1/NIMIN2-binding domain regions of Arabidopsis NPR1 to NPR4 in yeast. Sequences harboring LENRV-like motifs were expressed in fusion with the sequence for the Gal4 DNA-binding domain (BD), and sequences encoding N1/N2BDs were expressed in fusion with the sequence for the Gal4 transactivation domain (AD). Quantitative Y2H assays were conducted in absence and presence of salicylic acid. (a) Interaction between separated LEKRV and N1/N2BD regions from NPR3 and NPR4. BD–NtNPR1-LENRV/AD–NtNPR1-N1/N2BD interaction served as a positive control (Neeley *et al*., 2019). (b) Interaction between separated LEKRV and N1/N2BD regions of NPR3. Assays were conducted with cells grown without addition of chemicals or with cells grown in medium supplemented with SA or 4-hydroxybenzoic acid (4-OH BA). (c) Chimeric interactions between separated LENRV-like and N1/N2BD regions of NPR1 to NPR4.

Inability of association between NPR1 and NPR2 LENRV-like and N1/N2BD parts, respectively, may suggest that binding of SA is too weak or that the LENRV-like and N1/N2BD regions of these proteins are not able for steric reasons to associate a stable SA-binding fold. To analyze whether boosted positive charge could force C-terminal regions into physical contact, we changed asparagine 431 to lysine in NPR1 yielding the sequence LEKRV as found in NPR3 and NPR4 proteins that support SA-mediated remodeling of their C-terminal regions. However, mutant NPR1-LEKRV did not exhibit activity in the Y2H association assay (data not shown). Furthermore, by performing hybrid interaction assays, we tested whether failure to dimerize NPR1 and NPR2 C-terminal parts could be attributed to just one half of the SA fold. Indeed, we found that NPR1 LENRV part did not support association with any N1/N2BD region, while NPR1 N1/N2BD part displayed dimerization with LENRV-like regions from all NPR2, NPR3, NPR4 and even from NtNPR1, in an SA-dependent fashion (Fig. 3c and data not shown). Conversely, the N1/N2BD part of NPR2 was completely inactive, while NPR2 YENRV part was able to bind N1/N2BD regions from NPR1, NPR3 and NPR4 (Fig. 3c). Thus, the LENRV region of NPR1 and the N1/N2BD region of NPR2 occlude SA-induced intramolecular remodeling in the C-termini of these proteins.

### The conserved arginine residue in the LENRV-like motifs mediates salicylic acid responsiveness of all NPR1 to NPR4

As shown in our previous work, binding of NIMIN1 and NIMIN2 proteins to NPR1 is clearly inhibited in presence of SA, showing that NIMIN1/NIMIN2 interaction renders NPR1 sensitive to SA, and we have already demonstrated that the arginine residues in the LENRV motifs of Arabidopsis and tobacco NPR1 (R432 in AtNPR1; R431 in NtNPR1) are critical for SA-induced relief from NIMIN1/NIMIN2 binding (Maier *et al*., 2011). Furthermore, R431 supports both transcription enhancement and interaction of C-terminal LENRV and N1/N2BD regions in NtNPR1 (Maier *et al*., 2011; Neeley *et al*., 2019), and AtNPR4 binds SA via R419 (Ding *et al*., 2018; Wang *et al*., 2020). Since Arabidopsis NPR1 to NPR4 all harbor arginine at position 4 in their LENRV-like motifs (Fig. S7), we mutagenized the residue to lysine, as found in the inactive *nim1-4* mutant (Ryals *et al*., 1997; Canet *et al*., 2010) and monitored protein-binding activities of NPR2, NPR3 and NPR4 mutant proteins in absence and presence of SA.

For NPR2, we changed R432 to lysine in context of the full-length protein and studied interaction of mutant NPR2 with TGA7 (for an overview of mutants described in this section, see Fig. S1c). NPR2 R432K is still able to bind TGA7 as does wild-type protein. However, SA-induced enhancement of binding is completely lost (Fig. 4a). Similarly, mutation of R428 in NPR3 and R419 in NPR4, yielding the sequence LEKKV, prevented interaction of C-terminal NPR3 and NPR4 domains, respectively (Fig. 4b,c), while mutation of K418 to arginine in NPR4 (LERRV) did not impair SA-dependent binding of separated NPR4 parts (Fig. 4c). On the other hand, switching lysine and arginine residues at positions 418/419 in NPR4 (LERKV) could not rescue the LEKKV mutant (Fig. 4c). Together, the data substantiate the unique and general significance of arginine embedded in a LENRV-like signature for SA sensitivity of all NPR1 to NPR4 and corroborate the strategic position of R419 in NPR4 for ligand binding deduced from X-ray crystallography (Wang *et al*., 2020).

**Fig. 4.**
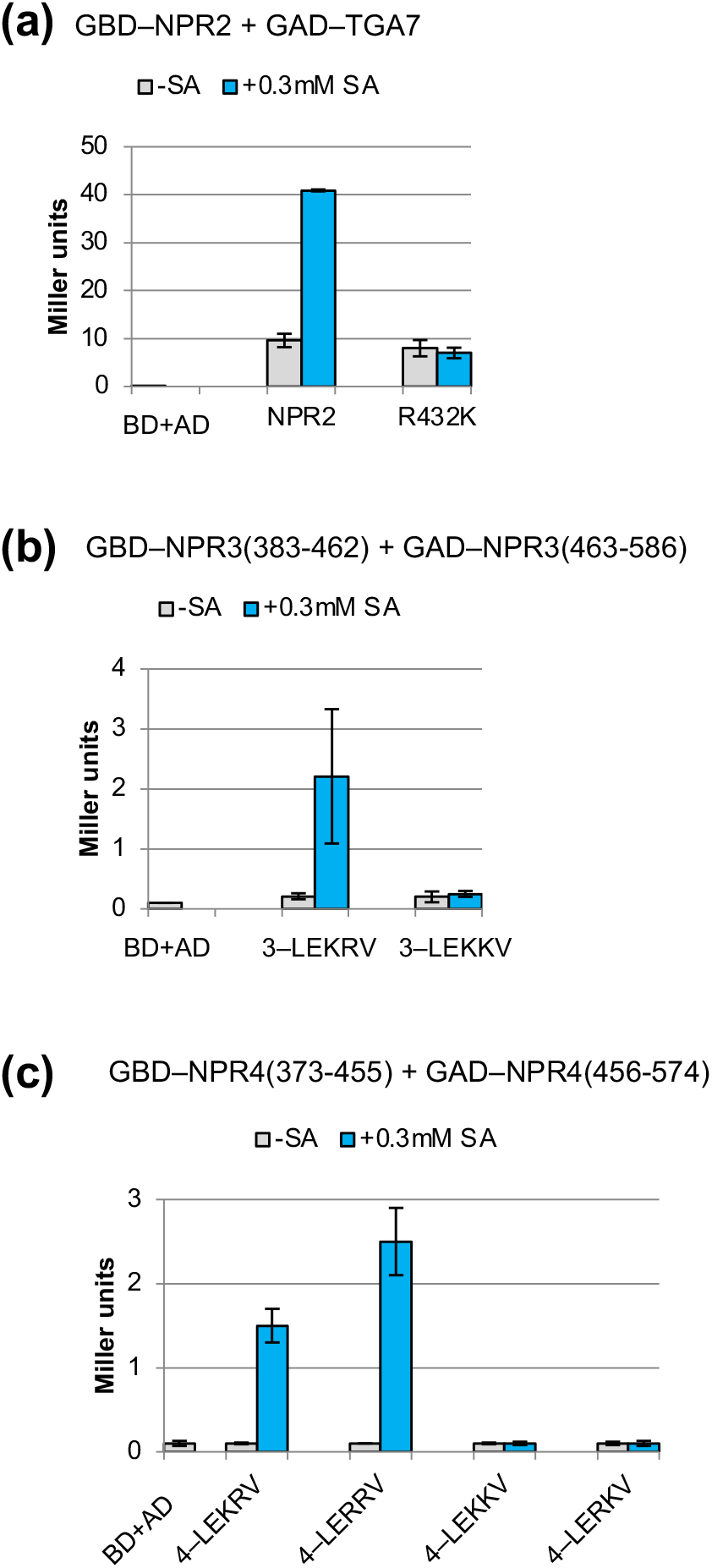
Effect of mutations in the LENRV-like motif on protein-protein interactions of Arabidopsis NPR2 to NPR4. Sequences harboring the LENRV-like motifs of NPR2 to NPR4 were expressed in fusion with the sequence for Gal4 DNA-binding domain (BD), and sequences encoding N1/N2BDs and TGA7 were expressed in fusion with the sequence for Gal4 transactivation domain (AD). Quantitative Y2H assays were conducted in absence and presence of salicylic acid. Activities of mutants are compared to acitity of the respective wild-type protein. (a) Interaction of NPR2 R432K with TGA7. (b) Interaction of NPR3(383-462) R428K with NPR3(463-586). (c) Interaction of NPR4(373-455) mutants K418R, R419K and double mutant K418R R419K with NPR4(456-574).

### NPR1 to NPR4 display differential sensitivity to SA

All Arabidopsis NPR1 to NPR4 respond to SA. While NPR3 and NPR4 are directly accessible to SA associating their LEKRV and N1/N2BD parts in an SA-dependent fashion, NPR1 and NPR2 cannot perform this reaction. However, association with interaction partners, NIMIN1/NIMIN2 and TGA7, respectively, renders NPR1 and NPR2 accessible to SA perception. To further elucidate the sensitivity of signal perception, we determined half-maximal SA concentrations affecting the different protein-protein interactions of Arabidopsis NPR1 to NPR4.

For NIMIN2-NPR1 interaction, we measured a half-maximal inhibiting SA concentration (IC_50_) of 17µM (Fig. 5a). This concentration is one order higher than the value for inhibition of NtNIMIN2a-NtNPR1 interaction (IC_50_ = 0.7µM; Maier *et al*., 2011; Neeley *et al*., 2019) indicating that AtNPR1 is less sensitive to SA than NtNPR1. The half-maximal enhancing concentration (EC_50_) for TGA7-NPR2 interaction was in the same range as for inhibition of NIMIN2-NPR1 interaction (EC_50_ = 19µM; Fig. 5b). Yet, association of NPR3 and NPR4 LEKRV and N1/N2BD parts, respectively, by SA could not be saturated under our assay conditions implying that NPR3 and NPR4 are considerably less sensitive to SA than NPR1 and NPR2 in complex with their respective interaction partners (NPR3 EC_50_ > 121µM; NPR4 EC_50_ > 67µM; Fig. 5c,d).

**Fig. 5.**
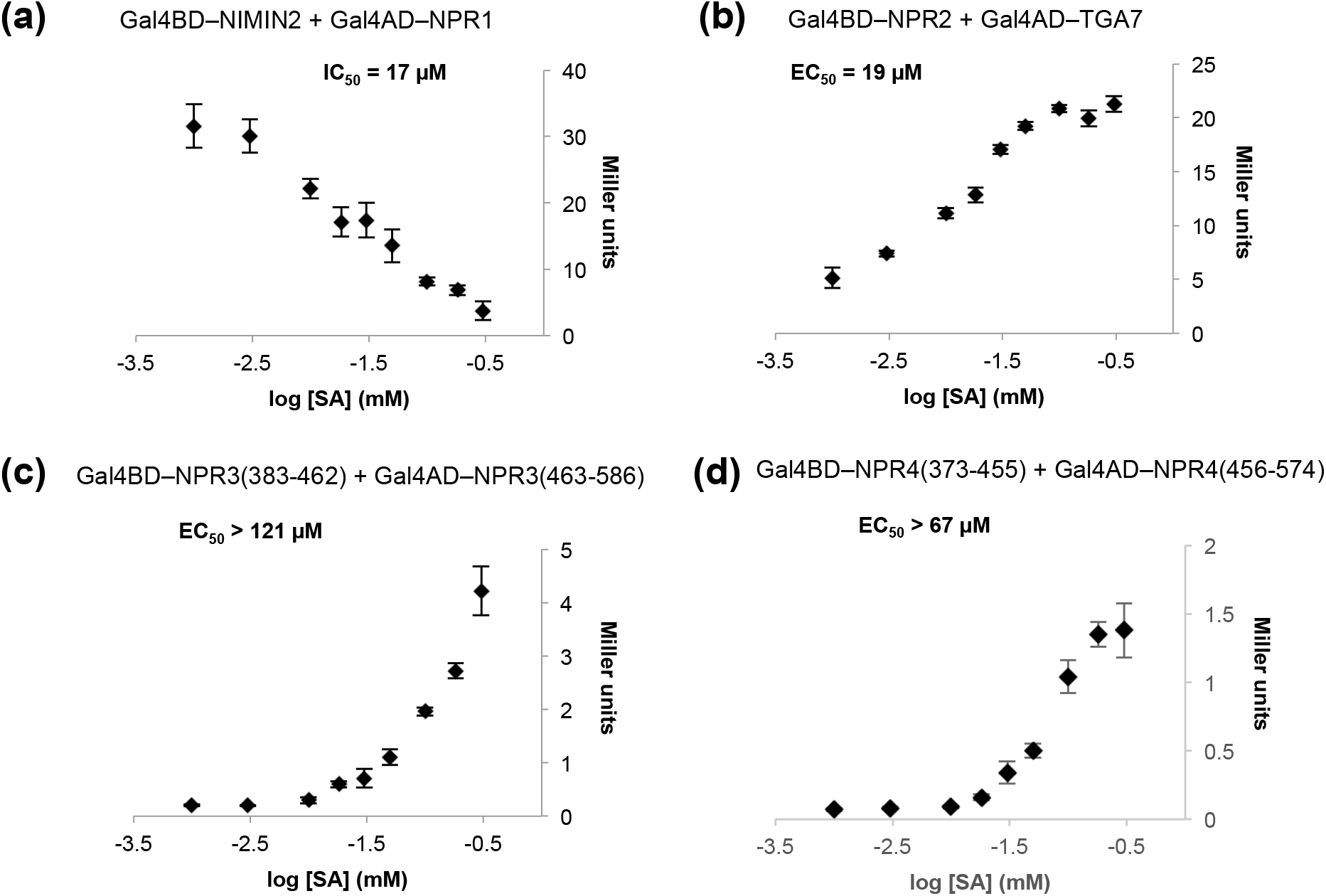
Determination of half-maximal salicylic acid concentrations affecting protein-protein interactions of Arabidopsis NPR1 to NPR4 in yeast. (a) Interaction of NIMIN2 with NPR1. (b) Interaction of NPR2 with TGA7. (c) Interaction of NPR3(383-462) with NPR3(463-586). (d) Interaction of NPR4(373-455) with NPR4(456-574).

### NPR1 to NPR4 display differential sensitivity to benzoic acid derivatives

We also tested the impact of synthetic SA analogs on NPR protein-protein interactions. Several structural analogs of SA have been shown to induce *PR* genes and disease resistance *in planta* (Métraux *et al*., 1991; Conrath *et al*., 1995; Görlach *et al*., 1996; Friedrich *et al*., 1996; Knoth *et al*., 2009; Palmer *et al*., 2019), and we have already demonstrated that functional analogs BTH and INA and some benzoic acid derivatives can replace SA in YH analyses of NtNPR1 (Neeley *et al*., 2019). Here, we tested SA analogs (Fig. S9) respecting their ability to induce activity changes of Arabidopsis NPR1 to NPR4. We considered NIMIN2 interaction for NPR1, TGA7 interaction for NPR2, and for NPR3 and NPR4, we examined association of separated C-terminal domains LEKRV and N1/N2BD.

Among SA, BTH and INA, SA was the most potent agent acting on NPR1, NPR3 and NPR4 (Fig. S10), while acetylsalicylic acid and 5-fluorosalicylic acid, on the other hand, induced similar activities as SA (Fig. 6a-c). However, one chemical made a clear difference. 3,5-Dichloroanthranilic acid (3,5-DCA, DCA) proved more effective on NPR1 than SA (Fig. 6a, Fig. S11a). NIMIN2-NPR1 complex was about four times more sensitive to DCA than to SA (IC_50(DCA)_ = 4µM; Fig. 6d). Effects were even more pronounced with association of NPR3 C-terminal domains being at least twenty times more sensitive to DCA than to SA (EC_50(DCA)_ = 6µM; Fig. 6b,e, Fig. S11b). Surprisingly, association of NPR4 C-terminal domains, in contrast to NPR3 domains, was essentially unresponsive to DCA (Fig. 6c, Fig. S11c).

**Fig. 6.**
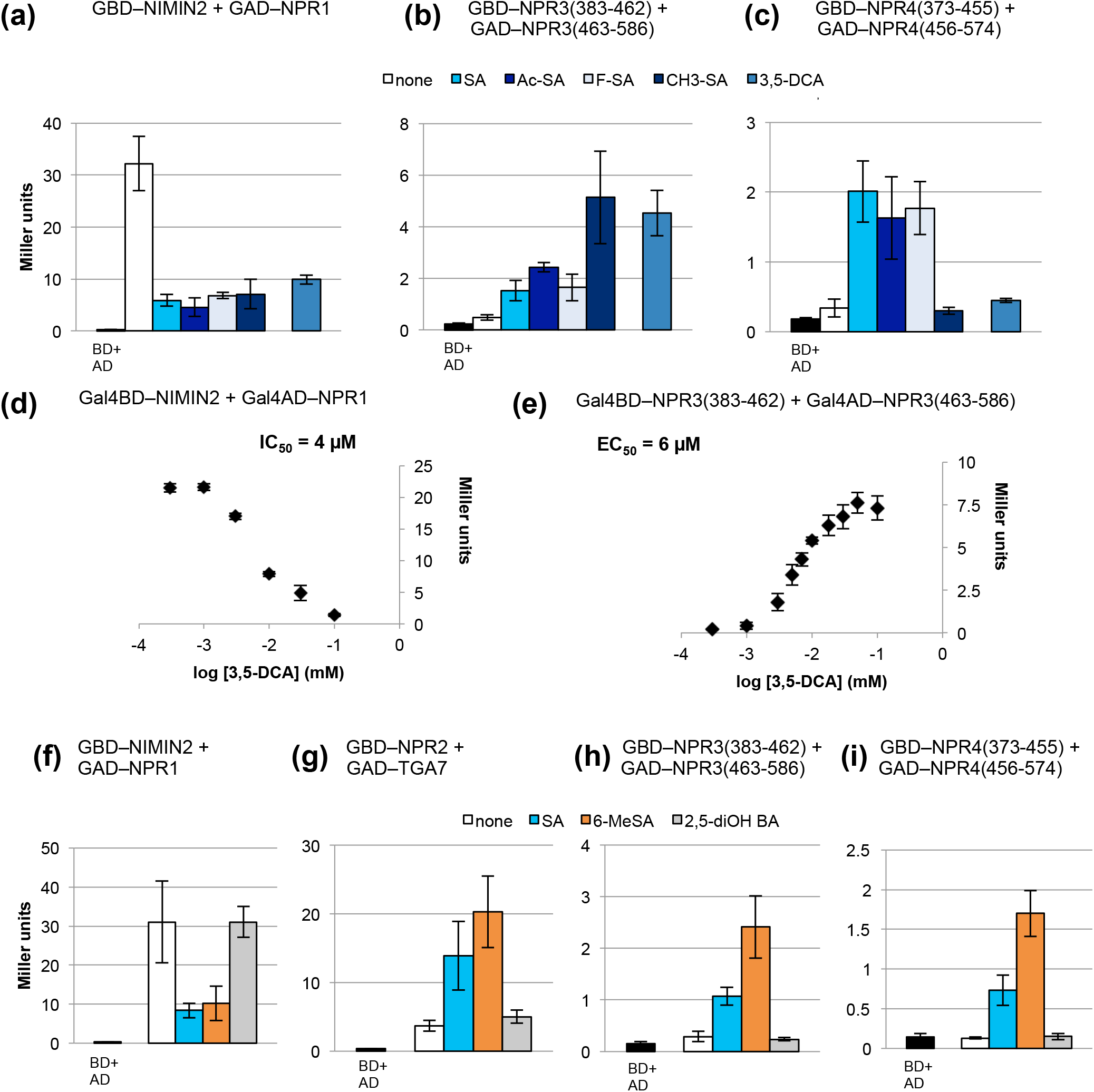
Effects of salicylic acid analogs on protein-protein interactions of Arabidopsis NPR1 to NPR4 in yeast. Quantitative Y2H assays were conducted with cells grown without addition of chemicals or with cells grown in medium supplemented with salicylic acid or analogs. Concentration of chemicals was 300µM apart from 3,5-DCA (30µM). Chemicals were solved in DMSO. The final DMSO concentration in yeast growth medium was 0.1%. (a,f) Interaction of NIMIN2 with NPR1. (b,h) Interaction of NPR3(383-462) with NPR3(463-586). (c,i) Interaction of NPR4(373-455) with NPR4(456-574). (d) Determination of the half-maximal 3,5-DCA concentration for NIMIN2-NPR1 interaction. (e) Determination of the half-maximal 3,5-DCA concentration for NPR3(383-462)-NPR3(463-586) interaction. (g) Interaction of NPR2 with TGA7. SA, salicylic acid; Ac-SA, acetylsalicylic acid; F-SA, 5-fluorosalicylic acid; CH3-SA, 5-methylsalicylic acid; 3,5-DCA, 3,5-dichloroanthranilic acid; 6-MeSA, 6-methylsalicylic acid; 2,5-diOH BA, 2,5-dihydroxybenzoic acid.

### Both the extended LEKRV and the N1/N2BD regions determine specificity of ligand binding to the NPR4 C-terminus

Unresponsiveness of NPR4 C-terminal domains to some structural SA analogs suggested that the NPR4 binding pocket enclosing the SA molecule is tailored rather tight, while NPR3 and NPR1 are able to perceive compounds with bulky substituents on the hydrophobic benzene ring. Mutagenesis had shown that the arginine residue in the LENRV-like motifs of NPR1 to NPR4 is indispensable for hormone sensing, and that, at least for NPR4, position of the arginine within the LEKRV motif is critical.

To analyze the contributions of the LEKRV extended region and the N1/N2BD to ligand binding, we tested chimeric protein-protein interactions using the NPR4-LEKRV part in combination with N1/N2BD regions from NPR4, NPR3 and NPR1, and the NPR4 N1/N2BD together with the NPR3-LEKRV region. Reconstitution of ligand-binding folds by SA was observed with all combinations in a similar fashion. Yet, NPR3-N1/N2BD part shifted ligand selection of the NPR4-NPR3 chimera towards DCA, while NPR1-N1/N2BD part supported an even more promiscuous hybrid receptor accepting various agonists (Fig. S12a-c). Similarly, using the NPR3-LEKRV region in combination with NPR4-N1/N2BD part fostered accessibility of DCA to the hormone binding sites (Fig. S12d). Together, the data are consistent with the X-ray crystallographic data demonstrating that both the LEKRV part and the N1/N2BD participate in forming the ligand-binding pocket of NPR4.

### NPR1 to NPR4 respond to a microbial salicylic acid derivative

In addition to synthetic analogs, we tested two SA derivatives occurring naturally in plants and microbes, respectively. Gentisic acid (GA; 2,5-dihydroxybenzoic acid [2,5-diOH BA]) is a secondary metabolite found in many plants and fruits, like gentian, grapes, citrus fruits, kiwi and blackberries (Griffiths, 1959; Abedi *et al*., 2020). It is a major catabolite which is synthesized from SA by hydroxylation (Zhang *et al*., 2017). GA has also been shown to be induced in viroid and virus-infected tomato leading to accumulation of a subset of PR proteins different from PR-1 (Bellés *et al*., 1999). With GA, we were not able to detect effects on reporter activity in NPR Y2H interactions (Fig. 6f-i).

6-Methylsalicylic acid (6-MeSA) is a microbial metabolite. It is widely present in actinomycetes, fungi and lichens where it is a building block for many bioactive secondary products, f.e., the *Penicillium* toxin patulin and the *Streptomyces* antibiotic cholorothricin (Ding *et al*., 2010). The compound is synthesized by 6-MeSA synthase enzymes sharing high homologies in sequence and domain organization in fungi and bacteria. Interestingly, 6-MeSA exerted activity in all Y2H combinations we tested. Regarding effects on NIMIN2-NPR1 and NPR2-TGA7 interactions, 6-MeSA was about equally active as SA, but with NPR3 and NPR4 LEKRV-NBD associations, 6-MeSA was about double active compared to SA (Fig. 6f-i, Fig. S13a,b). The EC_50(6-MeSA)_ value for NPR3 LEKRV-N1/2BD association was 151µM (Fig. S13c).

### Response of SAR gene promoters to SA analogs

To review binding of different SA analogs to NPR1, we tested chemical activation of SA-inducible promoters using transgenic tobacco lines harboring *-1533PR-1a:GUS* and *NIMIN1:GUS* reporter gene constructs (Grüner *et al*., 1994; Glocova *et al*., 2005). SA-mediated transcription of both *PR-1* and *NIMIN1* is dependent on NPR1 (Ryals *et al*., 1997; Shah *et al*., 1997; Hermann *et al*., 2013). In consent with a previous report (Palmer *et al*., 2019), we observed strong induction of the two promoters by acetylsalicylic acid and 5-fluorosalicylic acid (Fig. S14), presumably reflecting the potential of these analogs to bind to NPR1. With 6-methylsalicylic acid, we detected moderate activation of the two promoters.

These latter findings corroborate a previous report on induction of PR-1 proteins by treatment of tobacco plants with 6-MeSA (Yalpani *et al*., 2001). We were not able, however, to detect significant reporter enzyme activity with 2,5-dihydroxybenzoic acid, which does not bind to NPR1, and with 0.1mM DCA (Fig. S14).

## Discussion

NPR1 is the explicit positive regulator of SA-induced *PR-1* gene expression and SAR in Arabidopsis. The gene is part of a small family comprising four members, NPR1 to NPR4, harboring the two C-terminal domains LEKRV-like and N1/N2BD involved in binding the SAR signal SA in tobacco NPR1 and Arabidopsis NPR4. NPR3 and NPR4 have also been implied in *PR-1* gene regulation, although different models have been launched. Indeed, genetic, biochemical and structural evidence has demonstrated that NPR1 to NPR4 are targeted by the SA signal. Here we compared Arabidopsis NPR1 to NPR4 in a biochemical approach using various YH assays to explore the functional potential of NPR proteins. Table 1 summarizes our results.

**Table 1.** Biochemical capabilities of Arabidopsis NPR1 to NPR4 as determined by yeast hybrid analyses.

| Activity | NPR1 | NPR2 | NPR3 | NPR4 |
| --- | --- | --- | --- | --- |
| transcription activity | no | no | no | no |
| LENRV-like + N1/2BD association | no | no | EC<sub>50</sub>>121 µM | EC<sub>50</sub>>67 µM |
| NIMIN1/NIMIN2 interaction | NIMIN1<br>N2: IC<sub>50</sub> = 17 µM | no | no | no |
| NIMIN3 interaction | ✓ | no | no | no |
| TGA factor interaction | ✓ | TGA7:<br>EC<sub>50</sub> = 19 µM | ✓ | ✓ |
| formation of ternary NIMIN–NPR–TGA complexes | N2, N3 ✓ | n.d. | no | no |
|  | N1 |  |  |  |
| SA sensitivity dependent on R in the LENRV-like motif | ✓ | ✓ | ✓ | ✓ |
| sensitivity towards 3,5-DCA | N2: IC<sub>50</sub> = 4 µM | n.d. | LEKRV+NBD:<br>EC<sub>50</sub> = 6 µM | no |
| sensitivity towards 6-MeSA | NIMIN2 | TGA7 | LEKRV+NBD | LEKRV+NBD |

3,5-DCA, dichloroanthranilic acid; 6-MeSA, 6-methylsalicylic acid; N, NIMIN protein; NBD, NIMIN1/NIMIN2-binding domain
activity
activity affected by SA (>3-fold)
activity affected by SA analog (>3-fold)
no activity
n.d. not determined

### Yeast hybrid assays as a tool to study salicylic acid signaling through NPR proteins

Based on YH assays (Maier *et al*., 2011; Neeley *et al*., 2019) and in accordance with genetic evidence (Ryals *et al*., 1997; Canet *et al*., 2010), we were able to devise a mechanism for perception of the SAR signal SA by tobacco NPR1 in the past. SA was predicted to bind via its carboxyl to the guanidinium group of the arginine in the penta-amino acid motif LENRV and to induce physical association of the two conserved domains LENRV and N1/N2BD. Furthermore, association of two distinct domains in the NPR1 C-terminus was anticipated to induce a conformational switch, which, in tobacco NPR1, coincides with SA-mediated transcription enhancement and binding of TGA transcription factors to the C-terminus (Neeley *et al*., 2019). Essentially, this mechanism of SA perception has recently been confirmed by X-ray crystallography for Arabidopsis NPR4 harboring conserved LEKRV and N1/N2BD stretches which both contact the SA molecule (Wang *et al*., 2020). Together, our results demonstrate that YH analyses are an appropriate tool to gain mechanistic insight into signaling through NPR family members.

### Basal mechanism of SA perception by Arabidopsis NPR1 to NPR4

In extension of current knowledge, we demonstrate that NPR1 to NPR4 are sensitive to the SA signal in yeast, and that SA sensitivity of all NPR1 to NPR4 relies on the arginine residue embedded in the penta-amino acid motifs LENRV in NPR1, YENRV in NPR2 and LEKRV in NPR3 and NPR4. Both NPR3 and NPR4 display association of their C-terminal LEKRV and N1/N2BD regions in an SA-dependent manner in yeast, thus corroborating previous results obtained with tobacco NPR1 (Neeley *et al*., 2019) and the X-ray crystallographic data for SA-NPR4 complex (Wang *et al*., 2020). Hence, all NPR1 to NPR4 are receptors for the SA signal. Recent genetic evidence has come to the same conclusion (Castelló *et al*., 2018).

Quite surprisingly, however, neither NPR2 nor NPR3 nor NPR4 could bind NIMIN1 or NIMIN2 in Y2H assays, although the proteins harbor a highly conserved stretch in their C-termini that, through mutagenesis, was identified as binding site for NIMIN2-type proteins in tobacco NPR1 (Maier *et al*., 2011; Neeley *et al*., 2019). As a matter of fact, the N1/N2BD region of NPR4, as defined by us (aa 484 to 500), falls together with the cluster of seven amino acid residues from positions 488 to 503 forming part of the SA-binding pocket (Fig. S7; Wang *et al*., 2020). Thus, the conserved C-terminal domain, which we identified as N1/N2BD in tobacco NPR1 and which interacts with the LENRV part of NtNPR1 in an SA-dependent fashion, serves primarily as SA-binding interface. Consequently, NPR1 to NPR4 likely perceive the SA signal via the same basal mechanism engaging the arginine in the LENRV-like motif, the extended LENRV-like region and the N1/N2BD for trapping the SA molecule to achieve protein activation (Wang *et al*., 2020). Yet, we infer that the domain termed N1/N2BD by us, or part of it, is also involved in interaction with NIMIN1 and NIMIN2 proteins in tobacco and Arabidopsis NPR1. In fact, we have already shown that NIMIN1- and NIMIN2-type proteins can bridge separated LENRV and N1/N2BD parts of NtNPR1 (Neeley *et al*., 2019). Therefore, and for the sake of clarity and simplicity, we will further allude to this conserved amino acid stretch as N1/N2BD even though it is not clear whether NPR2 to NPR4 indeed bind NIMIN1 or NIMIN2 via this region (Fig. S7). Of note, NPR1 homologs from all higher plant species harbor the two conserved domains, LEKRV-like motif with the arginine at position 4 and N1/N2BD, in their C-termini.

### Differential sensitivity of Arabidopsis NPR1 to NPR4 to the SA signal

Although all NPR1 to NPR4 likely employ the same basal mechanism for SA perception, there are nevertheless major differences between them. Most importantly, both clade 1 proteins, NPR1 and NPR2, cannot associate their LENRV-like and N1/N2BD regions in presence of SA in yeast suggesting that they cannot perceive the SA signal on their own. Our findings corroborate previous *in vitro* assays demonstrating that NPR1, in contrast to NPR3 and NPR4, cannot bind SA (Fu *et al*., 2012; Wang *et al*., 2020). Rather, both clade 1 proteins NPR1 and NPR2 require interaction with partners enabling them to sense SA. Thus, NIMIN1/NIMIN2 and some TGA factors, e.g., TGA7, function as co-factors in SA signal transduction through NPR1 and NPR2, respectively. In this line, binding of NIMIN1 or NIMIN2 at or close to the SA contact regions would shape the NPR1 C-terminal region to facilitate hormone sensing.

In contrast to NPR1 and NPR2, clade 2 proteins NPR3 and NPR4 can bind the SA signal directly. Although we were not able to obtain precise EC_50_ values for SA binding to NPR3 and NPR4, it was evident that, in agreement with other reports (Fu *et al*., 2012; Ding *et al*., 2018), NPR3 is less sensitive to SA than NPR4. This finding was supported by a less stringent ligand binding specificity of NPR3 compared to NPR4. In contrast to NPR4, NPR3 is able to accommodate ligands with halogen substituents at positions 3 and 5 on the benzene ring, implying that NPR3´s affinity for non-substituted SA is weaker. Our findings on differential affinities of NPR1 to NPR4 to the SA molecule (NPR1, NPR2 << NPR3 < NPR4), determined through Y2H analyses, are in full accord with previous *in vitro* SA binding assays and with X-ray crystallographic data of SA bound to NPR4 and to NIMIN2-NPR1 complex (Fu *et al*., 2012; Wang *et al*., 2020).

### Function of NPR1 as SA receptor

All NPR1 to NPR4 can perceive the SA signal. However, genetic analyses have shown unambiguously that NPR1 is the exclusive positive regulator of *PR-1* gene expression in response to SA in Arabidopsis that cannot be replaced by the other NPR proteins (Liu *et al*., 2005; Zhang *et al*., 2006; Castelló *et al*., 2018). Using a biochemical approach, we demonstrate that NPR1 stands out from the paralogs by two unique features. First, NIMIN1 and NIMIN2 bind strongly to the NPR1 C-terminus, and second, binding of NIMIN1 and NIMIN2 proteins renders NPR1 sensitive to SA. Together, these features are likely instrumental for the role of NPR1 as SAR regulator. In fact, it came as a big surprise that NPR1 cannot perceive the SA signal on its own, but rather requires NIMIN1/NIMIN2 binding to enable SA sensing. Of note, *NIMIN1* and *NIMIN2* are not expressed constitutively (Weigel *et al*., 2001, 2005). Both genes are induced themselves by SA prior to *PR-1* gene induction (Hermann *et al*., 2013). Altogether, sensitizing NPR1 to SA through interaction with SA-induced NIMIN1 or NIMIN2 is in accord with the previous finding that transcription of the tobacco *PR-1a* gene requires *de novo* protein synthesis (Qin *et al*., 1994). The mechanism ensures that NPR1 activity is strictly controlled and can occur only after a certain threshold level of SA has been exceeded. Consistently, it has been shown that *PR-1* transcripts and PR-1 proteins accumulate late after exogenous chemical induction, in pathogen-infected plant tissue and in the course of SAR (Ward *et al*., 1991; Horvath *et al*., 1998; Weigel *et al*., 2005; Hermann *et al*., 2013). A threshold-dependent induction of *PR-1* delays protein synthesis to some extent, but, in the end, secures strong and robust PR-1 accumulation in tissue substantially threatened by pathogen attack. Such mechanism of *PR-1* gene regulation is also well in line with the phenomenon of priming (Hermann *et al*., 2013). In priming, a low level SA stimulus is initially not translated into active defense. However, faster and/or stronger accumulation of PR-1 proteins is observed in response to SA challenge doses (Conrath *et al*., 2002; Mauch-Mani *et al*., 2017). On the other hand, undue protein accumulation, as experienced in *NIMIN1* overexpression in Arabidopsis and in *NIMIN2a* overexpression in tobacco plants (Weigel *et al*., 2005; Zwicker *et al*., 2007), is likely to compromise SA perception via NPR1. Thus, basic insensitivity of constitutively accumulating NPR1 to low levels of SA and competition of SA and NIMIN1/NIMIN2 for binding to the NPR1 C-terminus seem critical aspects of well-balanced and likewise effective pathogen defense conveyed through NPR1-dependent PR proteins in non-infected systemic leaves. Altogether, the mechanism envisaged by us for activation of NPR1 (Fig. 7) meets both specificity on the one hand and physiological features associated with SA-induction of *PR-1* on the other hand.

**Fig. 7.**
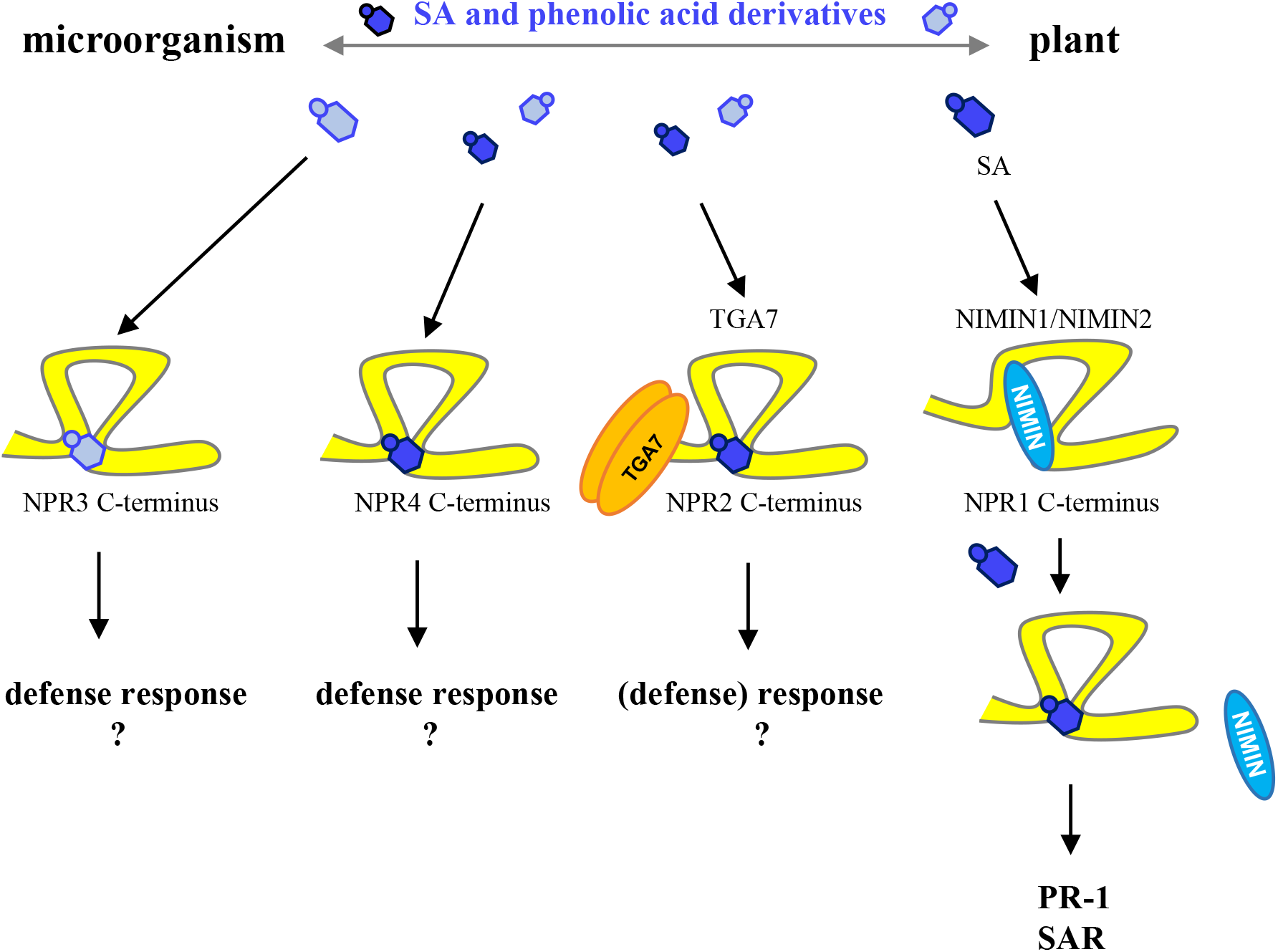
Working model for differential activities of the salicylic acid receptors NPR1 to NPR4 in Arabidopsis. Interaction of plants with microbes produces salicylic acid and other phenolic acid derivatives. Although NPR1 to NPR4 perceive SA and structural analogs by the same basal mechanism, the receptors activate distinct pathogen defense responses that proceed in parallel. NPR1 is the canonical SAR receptor activated by SA which is synthesized by the plant after microbial attack. Sensitizing NPR1 for SA requires prior interaction with SA-induced NIMIN1 or NIMIN2. In contrast, NPR3 and NPR4 are activated directly by the signal. NPR4 prefers SA, while NPR3 likely binds an unknown SA analog which may be of microbial or plant origin. Finally, NPR2, like NPR1, needs interaction with a co-factor, f.e., TGA7, to enable perception of the SA signal. Differential interaction of the receptors of the NPR family with SA, SA analogs and co-factors facilitating signal perception are a likely biochemical basis for distinct signaling outputs of NPR1 to NPR4.

### Functional significance of NPR3

Although both clade 2 proteins NPR3 and NPR4 bind SA directly, there are nevertheless clear differences between them regarding affinity for SA and specificity of ligand binding.

Consistent with the X-ray crystallographic data, implying that the NPR4 C-terminus buries the SA molecule in a cavity by engaging 14 amino acid residues to form the SA binding pocket (Wang *et al*., 2020), we demonstrate that NPR4 is rather selective for the SA ligand. In contrast, NPR3 displays low sensitivity to SA and much higher sensitivity to DCA which is not bound by NPR4 suggesting different spatial organization of the respective ligand binding regions. Indeed, we show that ligand selection can be modified by combining NPR4 and NPR3 or NPR1 C-terminal parts in hybrid receptors indicating that the C-terminal regions of NPR3 and NPR4 are molded by three participants, the ligand, the LENRV region and the N1/N2BD.

Distinct ligand preferences of NPR3 and NPR4 are indeed striking as it was suggested in one model that NPR3 and NPR4 act redundantly as transcriptional co-repressors of defense gene expression and that repression is released by SA (Ding *et al*., 2018). The NPR3/NPR4-mediated mechanism of *PR* gene repression and relief is unclear. Rather than underlining a redundant function for NPR3 and NPR4, our data would stress more distinct roles for the two receptors. DCA was identified in an elicitor screen for compounds inducing reporter gene expression in Arabidopsis from the promoter of a gene encoding a putative calmodulin-like calcium-binding protein (*CaBP22*; Knoth *et al*., 2009). The *CaBP22* promoter responds to several stimuli, including the oomycete *Hyaloperonospora parasitica*, the bacterial pathogen *Pseudomonas putida* and SA. However, in comparison to SA, DCA proved significantly more potent regarding *CaBP22* promoter activation and induced much stronger resistance against *H. parasitica* and *P. putida*. Importantly, DCA-triggered defense did not require accumulation of SA and was both NPR1-dependent and NPR1-independent. Furthermore, microarray analyses revealed that DCA, INA and BTH, which target NPR1 to NPR4 differentially in the Y2H assays conducted by us, induce both a common set of genes and unique transcriptional changes in Arabidopsis (Knoth *et al*., 2009; Bektas & Eulgem, 2015). Hence, in addition to NPR1, DCA signals through yet another unidentified receptor to elicit a subset of immune responses (Knoth *et al*., 2009). In this line, our results would support the notion that DCA, just like SA, targets both NPR1 and NPR3 to promote distinct pathogen defense responses in a dose-dependent manner that proceed in parallel. Similarly, mutant analysis revealed that perception of SA by NPR4 adds considerably to defense in Arabidopsis (Ding *et al*., 2018; Liu *et al*., 2020). Together, the data underscore differential roles for NPR3 and NPR4 acting as defense gene regulators independent from each other and from NPR1. Of note, such assumed function of NPR3 and NPR4 is also not well compatible with another model implying that NPR3 and NPR4, as parts of E3 ubiquitin ligase complexes, control availability of NPR1 by regulating the proteasomal degradation of NPR1 in an SA concentration-dependent manner (Fu *et al*., 2012).

Our finding of vastly differing sensitivity of NPR3 to DCA and SA suggests the possibility that NPR3 recognizes a signal molecule related to, but different from SA *in planta*. DCA is a synthetic compound not produced by plants. To explore the binding potential of NPR3 and its paralogs for natural SA analogs, we used SA derivatives known from plants and microbes. The plant-derived compound gentisic acid (2,5-dihydroxybenzoic acid) was inactive in the Y2H assays conducted by us and, consistent with previous studies (Abad *et al*., 1988), could not activate the NPR1-dependent *PR-1a* and *NIMIN1* promoters *in planta*.

However, 6-methylsalicylic acid, which is a common metabolite in fungi and bacteria and does not occur in plants, was sensed by all NPR1 to NPR4, with NPR3 and NPR4 displaying an even two-fold higher activity with 6-MeSA than with the SAR signal SA in yeast. The compound could also induce reporter gene expression from the *PR-1a* and *NIMIN1* promoters when applied to healthy tobacco leaf tissue. Thus, Arabidopsis NPR1 to NPR4 can bind a microbe-produced metabolite that impacts the receptors differentially. In this regard, it is interesting to note that defense responses were severely compromised in the Arabidopsis *npr1-1 npr4-4D* double mutant, even more dramatically than in the *SA-induction deficient2-1* (*sid2-1*) mutant (Liu *et al*., 2020) which is blocked in the prevailing route for microbe-induced SA biosynthesis in Arabidopsis (Garcion *et al*., 2008; Rekhter *et al*., 2019). This discrepancy has been attributed to residual levels of SA in *sid2-1*. However, NPR1-dependent *PR-1* gene expression was still severely compromised in the *sid2-1* mutant (Nawrath & Métraux, 1999; Liu *et al*., 2020). Together, our and other data are in consent with the notion that defense in Arabidopsis may not rely exclusively on the signal molecule SA. Rather, we suggest that the interaction of microorganisms with the plant delivers SA and compounds structurally related to SA. Such compounds could originate from different sources, not only from the plant, but also from microbes and/or different biosynthetic routes and could be bound preferentially, although not necessarily exclusively, by individual receptors of the NPR family. In particular, microbial signals may preferentially be perceived by clade 2 proteins NPR3 and NPR4 which are less sensitive to SA to activate distinct defense programs in parallel to NPR1. This then results in an additive and robust immune response depending on the flavor of the signals and tailored to best meet attack by heterogeneous microorganisms.

In this line, 6-MeSA, as a natural and common microbial metabolite that binds to NPR1 to NPR4, could serve as an external danger signal to activate defense in plants. Indeed, it has been demonstrated in a recent report that the nematophagous fungus *Duddingtonia flagrans* uses 6-MeSA, which is synthesized in hyphal tips, in three ways, as an intermediate for the biosynthesis of complex secondary compounds, as a morphogen to control formation of traps and as a chemoattractant to lure *Caenorhabditis elegans* worms into the fungal mycelium (Yu *et al*., 2021). Thus, in *D. flagrans*, 6-MeSA plays an important role in interkingdom communication between the fungus and nematodes (Yu *et al*., 2021).

## Conclusion

We show that Arabidopsis NPR family members NPR1 to NPR4 are *bona fide* SA receptors that perceive the signal via the same basal mechanism. Nevertheless, NPR1 to NPR4 clearly display unique biochemical qualities. We suggest that C-terminal domains LEKRV-like and N1/N2BD, in liaison with co-factors where applicable, facilitate perception of SA and structurally related molecules, enabling the plant to recognize attack by microbes and providing a biochemical basis for diverse defense signaling outputs activated independently and in parallel by NPR1 to NPR4 to ward off the attack. Figure 7 depicts our working model for defense signaling through the Arabidopsis NPR family of SA receptors.

## Acknowledgements

We thank David Neeley, Nadja Bluthardt, Ioanna Efstathiadou and Franziska Hadamjetz for YH analyses and Ingrid Prießnitz-Hohos for generation of transgenic plants. This work was supported by Land Baden-Württemberg and Universität Hohenheim.

## Author contributions

EK generated gene constructs, conducted most YH analyses and determined EC_50_ and IC_50_ values. MS performed immunodetections and YH assays. AJPP provided inestimable advice and fruitful discussions on the project. UMP generated gene constructs, was responsible for the coordination and supervision of the work, and wrote the article. All authors read and approved the manuscript.

## Supporting Information

Additional Supporting Information may be found online in the Supporting Information section at the end of the article.

**Fig. S1** Sequence alignment of C-terminal regions of Arabidopsis NPR1 to NPR4 and mutants used in Y2H analyses.

**Fig. S2** Accumulation of Arabidopsis NPR1 in leaf tissue.

**Fig. S3** Test for autoactivation of Arabidopsis NPR1 to NPR4 in yeast.

**Fig. S4** Test for homo- and heterodimerization among Arabidopsis NPR1 to NPR4 in yeast.

**Fig. S5** Interaction of Arabidopsis TGA transcription factors with truncated Arabidopsis NPR1 in yeast.

**Fig. S6** Formation of ternary complexes between Arabidopsis NIMIN2, TGA factors and NPR1, NPR3 or NPR4 in yeast.

**Fig. S7** Amino acid residues forming the NPR4 SA-binding pocket are conserved among Arabidopsis NPR1 to NPR4.

**Fig. S8** Yeast two-hybrid interaction between separated LENRV-like and NIMIN1/NIMIN2-binding domain regions of Arabidopsis NPR1 and NPR2, respectively.

**Fig. S9** Chemical structures of salicylic acid and analogs.

**Fig. S10** Effects of salicylic acid analogs on protein-protein interactions of Arabidopsis NPR1, NPR3 and NPR4 in yeast.

**Fig. S11** Effects of salicylic acid analogs on protein-protein interactions of Arabidopsis NPR1, NPR3 and NPR4 in yeast.

**Fig. S12** Formation of hybrid receptors by salicylic acid and structural analogs.

**Fig. S13** Effects of salicylic acid analogs on protein-protein interactions of Arabidopsis NPR3 and NPR4 in yeast.

**Fig. S14** Effects of salicylic acid analogs on *PR-1a* and *NIMIN1* promoter activity in transgenic tobacco plants.

**Table S1** Primers used for construction of full ORF and partial *NPR* cDNA clones.

**Table S2** Primers used for construction of mutant *NPR* cDNA clones.

**Table S3** Primers used for construction of TGA factor encoding cDNA clones.

**Methods S1** Detailed description of methods.

## References

Abad P, Marais A, Cardin L, Poupet A, Ponchet M. 1988. The effect of benzoic acid derivatives on Nicotiana tabacum growth in relation to PR-bl production. Antiviral Research 9: 315–327.

Abedi F, Razavi BM, Hosseinzadeh H. 2020. A review on gentisic acid as a plant derived phenolic acid and metabolite of aspirin: Comprehensive pharmacology, toxicology, and some pharmaceutical aspects. Phytotherapy Research 34: 729–741.

Altmann M, Altmann S, Rodriguez PA, Weller B, Vergara LE, Palme J, Marín-de la Rosa N, Sauer M, Wenig M, Villaécija JA et al. 2020. Extensive signal integration by the phytohormone protein network. Nature 583: 271–276.

Bektas Y, Eulgem T. 2015. Synthetic plant defense elicitors. Frontiers in Plant Science 5:804.

Bellés JM, Garro R, Fayos J, Navarro P, Primo J, Conejero V. 1999. Gentisic acid as a pathogen-inducible signal, additional to salicylic acid for activation of plant defenses in tomato. Molecular Plant-Microbe Interactions 12: 227–235.

Canet JV, Dobón A, Roig A, Tornero P. 2010. Structure-function analysis of npr1 alleles in Arabidopsis reveals a role for its paralogs in the perception of salicylic acid. Plant, Cell and Environment 33: 1911–1922.

Cao H, Bowling SA, Gordon AS, Dong X. 1994. Characterization of an Arabidopsis mutant that is nonresponsive to inducers of systemic acquired resistance. The Plant Cell 6: 1583–1592.

Cao H, Glazebrook J, Clarke JD, Volko S, Dong X. 1997. The Arabidopsis NPR1 gene that controls systemic acquired resistance encodes a novel protein containing ankyrin repeats. Cell 88: 57–63.

Castelló MJ, Medina-Puche L, Lamilla J, Tornero P. 2018. NPR1 paralogs of Arabidopsis and their role in salicylic acid perception. PLoS ONE 13(12):e0209835.

Conrath U, Chen Z, Ricigliano JR, Klessig DF. 1995. Two inducers of plant defense responses, 2,6-dichloroisonicotinic acid and salicylic acid, inhibit catalase activity in tobacco. Proceedings of the National Academy of Sciences USA 92: 7143–7147.

Conrath U, Pieterse CMJ, Mauch-Mani B. 2002. Priming in plant-pathogen interactions. TRENDS in Plant Science 7: 210–216.

Delaney TP, Friedrich L, Ryals JA. 1995. Arabidopsis signal transduction mutant defective in chemically and biologically induced disease resistance. Proceedings of the National Academy of Sciences USA 92: 6602–6606.

Dereeper A, Audic S, Claverie JM, Blanc G. 2010. BLAST-EXPLORER helps you building datasets for phylogenetic analysis. BMC Evolutionary Biology 10:8.

Després C, DeLong C, Glaze S, Liu E, Fobert PR. 2000. The Arabidopsis NPR1/NIM1 protein enhances the DNA binding activity of a subgroup of the TGA family of bZIP transcription factors. Plant Cell 12: 279–290.

Ding W, Lei C, He Q, Zhang Q, Bi Y, Liu W. 2010. Insights into bacterial 6-methylsalicylic acid synthase and its engineering to orsellinic acid synthase for spirotetronate generation. Chemistry & Biology 17: 495–503.

Ding Y, Sun T, Ao K, Peng Y, Zhang Y, Li X, Zhang Y. 2018. Opposite roles of salicylic acid receptors NPR1 and NPR3/NPR4 in transcriptional regulation of plant immunity. Cell 173: 1454–1467.

Friedrich L, Lawton K, Ruess W, Masner P, Specker N, Gut Rella M, Meier B, Dincher S, Staub T, Uknes S et al. 1996. A benzothiadiazole derivative induces systemic acquired resistance in tobacco. The Plant Journal 10: 61–70.

Fu ZQ, Dong X. 2013. Systemic acquired resistance: turning local infection into global defense. Annual Review of Plant Biology 64: 839–863.

Fu ZQ, Yan S, Saleh A, Wang W, Ruble J, Oka N, Mohan R, Spoel SH, Tada Y, Zheng N et al. 2012. NPR3 and NPR4 are receptors for the immune signal salicylic acid in plants. Nature 486: 228–232.

Garcion C, Lohmann A, Lamodière E, Catinot J, Buchala A, Doermann P, Métraux J-P. 2008. Characterization and biological function of the ISOCHORISMATE SYNTHASE2 gene of Arabidopsis. Plant Physiology 147: 1279–1287.

Glocova I, Thor K, Roth B, Babbick M, Pfitzner AJP, Pfitzner UM. 2005. Salicylic acid (SA)-dependent gene activation can be uncoupled from cell death-mediated gene activation: the SA-inducible NIMIN-1 and NIMIN-2 promoters, unlike the PR-1a promoter, do not respond to cell death signals in tobacco. Molecular Plant Pathology 6: 299–314.

Görlach J, Volrath S, Knauf-Beiter G, Hengy G, Beckhove U, Kogel KH, Oostendorp M, Staub T, Ward E, Kessmann H et al. 1996. Benzothiadiazole, a novel class of inducers of systemic acquired resistance, activates gene expression and disease resistance in wheat. The Plant Cell 8: 629–643.

Griffiths LA. 1959. On the distribution of gentisic acid in green plants. Journal of Experimental Botany 10: 437–442.

Grüner R, Pfitzner UM. 1994. The upstream region of the gene for the pathogenesis-related protein 1a from tobacco responds to environmental as well as to developmental signals in transgenic plants. European Journal of Biochemistry 220: 247–255.

Hermann M, Maier F, Masroor A, Hirth S, Pfitzner AJP, Pfitzner UM. 2013. The Arabidopsis NIMIN proteins affect NPR1 differentially. Frontiers in Plant Science 4:88.

Horvath DM, Huang DJ, Chua N-H. 1998. Four classes of salicylate-induced tobacco genes. Molecular Plant-Microbe Interactions 11: 895–905.

Klepikova AV, Kasianov AS, Gerasimov ES, Logacheva MD, Penin AA. 2016. A high resolution map of the Arabidopsis thaliana developmental transcriptome based on RNA-seq profiling. The Plant Journal 88: 1058–1070.

Knoth C, Salus MS, Girke T, Eulgem T. 2009. The synthetic elicitor 3,5-dichloroanthranilic acid induces NPR1-dependent and NPR1-independent mechanisms of disease resistance in Arabidopsis. Plant Physiology 150: 333–347.

Lebel E, Heifetz P, Thorne L, Uknes S, Ryals J, Ward E. 1998. Functional analysis of regulatory sequences controlling PR-1 gene expression in Arabidopsis. The Plant Journal 16: 223–233.

Liu G, Holub EB, Alonso JM, Ecker JR, Fobert PR. 2005. An Arabidopsis NPR1-like gene, NPR4, is required for disease resistance. The Plant Journal 41: 304–318.

Liu Y, Sun T, Sun Y, Zhang Y, Radojicic A, Ding Y, Tian H, Huang X, Lan J, Chen S et al. 2020. Diverse roles of the salicylic acid receptors NPR1 and NPR3/NPR4 in plant immunity. The Plant Cell 32: 4002–4016.

Maier F, Zwicker S, Hückelhoven A, Meissner M, Funk J, Pfitzner AJP, Pfitzner UM. 2011. NONEXPRESSOR OF PATHOGENESIS-RELATED PROTEINS1 (NPR1) and some NPR1-related proteins are sensitive to salicylic acid. Molecular Plant Pathology 12: 73–91.

Malamy J, Carr JP, Klessig DF, Raskin I. 1990. Salicylic acid: a likely endogenous signal in the resistance response of tobacco to viral infection. Science 250: 1002–1004.

Mauch-Mani B, Baccelli I, Luna E, Flors V. 2017. Defense priming: an adaptive part of induced resistance. Annual Review of Plant Biology 68: 485–512.

Métraux JP, Signer H, Ryals J, Ward E, Wyss-Benz M, Gaudin J, Raschdorf K, Schmid E, Blum W, Inverardi B. 1990. Increase in salicylic acid at the onset of systemic acquired resistance in cucumber. Science 250: 1004–1006.

Métraux JP, Ahl-Goy P, Staub T, Speich J, Steinemann A, Ryals J, Ward E. 1991. Induced systemic resistance in cucumber in response to 2,6-dichloroisonicotinic acid and pathogens. In: Hennecke H, Verma DPS, eds. Advances in molecular genetics of plant-microbe interactions. Dordrecht, Nl: Kluwer Academic Publishers, Vol. 1, 432–439.

Nawrath C, Métraux J-P. 1999. Salicylic acid induction–deficient mutants of Arabidopsis express PR-2 and PR-5 and accumulate high levels of camalexin after pathogen inoculation. The Plant Cell 11: 1393–1404.

Neeley D, Konopka E, Straub A, Maier F, Pfitzner AJP, Pfitzner UM. 2019. Salicylic acid-driven association of LENRV and NIMIN1/NIMIN2 binding domain regions in the C-terminus of tobacco NPR1 transduces SAR signal. Bioarchives. doi: 10.1101/543645

Niggeweg R, Thurow C, Weigel R, Pfitzner U, Gatz C. 2000. Tobacco TGA factors differ with respect to interaction with NPR1, activation potential and DNA-binding properties. Plant Molecular Biology 42: 775–788.

Palmer IA, Chen H, Chen J, Chang M, Li M, Liu F, Fu ZQ. 2019. Novel salicylic acid analogs induce a potent defense response in Arabidopsis. International Journal of Molecular Sciences 20:3356.

Qin X-F, Holuigue L, Horvath DM, Chua N-H. 1994. Immediate early transcription activation by salicylic acid via the cauliflower mosaic virus as-1 element. The Plant Cell 6: 863–874.

Rekhter D, Lüdke D, Ding Y, Feussner K, Zienkiewicz K, Lipka V, Wiermer M, Zhang Y, Feussner I. 2019. Isochorismate-derived biosynthesis of the plant stress hormone salicylic acid. Science 365: 498–502.

Rochon A, Boyle P, Wignes T, Fobert PR, Després C. 2006. The coactivator function of Arabidopsis NPR1 requires the core of its BTB/POZ domain and the oxidation of C-terminal cysteines. The Plant Cell 18: 3670–3685.

Ross AF. 1961. Systemic acquired resistance induced by localized virus infections in plants. Virology 14: 340–358.

Ryals J, Weymann K, Lawton K, Friedrich L, Ellis D, Steiner HY, Johnson J, Delaney TP, Jesse T, Vos P et al. 1997. The Arabidopsis NIM1 protein shows homology to the mammalian transcription factor inhibitor I kappa B. The Plant Cell 9: 425–439.

Shah J, Tsui F, Klessig DF. 1997. Characterization of a salicylic acid-insensitive mutant (sai1) of Arabidopsis thaliana, identified in a selective screen utilizing the SA-inducible expression of the tms2 gene. Molecular Plant-Microbe Interactions 10: 69–78.

Shi Z, Maximova S, Liu Y, Verica J, Guiltinan MJ. 2013. The salicylic acid receptor NPR3 is a negative regulator of the transcriptional defense response during early flower development in Arabidopsis. Molecular Plant 6: 802–816.

Stos-Zweifel V, Neeley D, Konopka E, Meissner M, Hermann M, Maier F, Häfner V, Pfitzner AJP, Pfitzner UM. 2018. Tobacco TGA7 mediates gene expression dependent and independent of salicylic acid. Bioarchives. doi: 10.1101/341834

Strompen G, Grüner R, Pfitzner UM. 1998. An as-1-like motif controls the level of expression of the gene for the pathogenesis-related protein 1a from tobacco. Plant Molecular Biology 37: 871–883.

van Loon LC, Rep M, Pieterse CM. 2006. Significance of inducible defense-related proteins in infected plants. Annual Review of Phytopathology 44: 135–162.

Vlot AC, Dempsey DA, Klessig DF. 2009. Salicylic acid, a multifaceted hormone to combat disease. Annual Review of Phytopathology 47: 177–206.

Wang W, Withers J, Li H, Zwack PJ, Rusnac D-M, Shi H, Liu L, Yan S, Hinds TR, Guttman M et al. 2020. Structural basis of salicylic acid perception by Arabidopsis NPR proteins. Nature 586: 311–316.

Ward ER, Uknes SJ, Williams SC, Dincher SS, Wiederhold DL, Alexander DC, Ahl-Goy P, Métraux J-P, Ryals JA. 1991. Coordinate gene activity in response to agents that induce systemic acquired resistance. The Plant Cell 3: 1085–1094.

Weigel RR, Bäuscher C, Pfitzner AJP, Pfitzner UM. 2001. NIMIN-1, NIMIN-2 and NIMIN-3, members of a novel family of proteins from Arabidopsis that interact with NPR1/NIM1, a key regulator of systemic acquired resistance in plants. Plant Molecular Biology 46: 143–160.

Weigel RR, Pfitzner UM, Gatz C. 2005. Interaction of NIMIN1 with NPR1 modulates PR gene expression in Arabidopsis. The Plant Cell 17: 1279–1291.

Yalpani N, Altier DJ, Barbour E, Cigan AL, Scelonge CJ. 2001. Production of 6-methylsalicylic acid by expression of a fungal polyketide synthase activates disease resistance in tobacco. The Plant Cell 13: 1401–1409.

Yu X, Hu X, Pop M, Wernet N, Kirschhöfer F, Brenner-Weiß G, Keller J, Bunzel M, Fischer R. 2021. Fatal attraction of Caenorhabditis elegans to predatory fungi through 6-methyl-salicylic acid. Nature Communications 12:5462.

Zavaliev R, Mohan R, Chen T, Dong X. 2020. Formation of NPR1 condensates promotes cell survival during the plant immune response. Cell 182: 1093–1108.

Zhang Y, Fan W, Kinkema M, Li X, Dong X. 1999. Interaction of NPR1 with basic leucine zipper protein transcription factors that bind sequences required for salicylic acid induction of the PR-1 gene. Proceedings of the National Academy of Sciences, USA 96: 6523–6528.

Zhang Y, Cheng YT, Qu N, Zhao Q, Bi D, Li X. 2006. Negative regulation of defense responses in Arabidopsis by two NPR1 paralogs. The Plant Journal: 48: 647–656.

Zhang Y, Zhao L, Zhao J, Li Y, Wang J, Guo R, Gan S, Liu CJ, Zhang K. 2017. S5H/DMR6 encodes a salicylic acid 5-hydroxylase that fine-tunes salicylic acid homeostasis. Plant Physiology 175: 1082–1093.

Zhou JM, Trifa Y, Silva H, Pontier D, Lam E, Shah J, Klessig DF. 2000. NPR1 differentially interacts with members of the TGA/OBF family of transcription factors that bind an element of the PR-1 gene required for induction by salicylic acid. Molecular Plant-Microbe Interactions 13: 191–202.

Zwicker S, Mast S, Stos V, Pfitzner AJP, Pfitzner UM. 2007. Tobacco NIMIN2 proteins control PR gene induction through transient repression early in systemic acquired resistance. Molecular Plant Pathology 8: 385–400.

