## Supporting Information for "Differential binding of salicylic acid, phenolic acid derivatives and co-factors determines the roles of Arabidopsis NPR1 to NPR4 in plant immunity"

**Evelyn Konopka, Mathias Saur, Artur J.P. Pfitzner and Ursula M. Pfitzner**
Fg. Allgemeine Virologie, Institut für Biologie, Universität Hohenheim, Emil-Wolff-Str. 14, D-70593 Stuttgart, Germany

ORCID ID Ursula M. Pfitzner: 0000-0002-7566-7614

*Author for correspondence:*
*Ursula Pfitzner*
**

**Fig. S1** Sequence alignment of C-terminal regions of Arabidopsis NPR1 to NPR4 and mutants used in Y2H analyses.

**Fig. S2** Accumulation of Arabidopsis NPR1 in leaf tissue.

**Fig. S3** Test for autoactivation of Arabidopsis NPR1 to NPR4 in yeast.

**Fig. S4** Test for homo- and heterodimerization among Arabidopsis NPR1 to NPR4 in yeast.

**Fig. S5** Interaction of Arabidopsis TGA transcription factors with truncated Arabidopsis NPR1 in yeast.

**Fig. S6** Formation of ternary complexes between Arabidopsis NIMIN2, TGA factors and NPR1, NPR3 or NPR4 in yeast.

**Fig. S7** Amino acid residues forming the NPR4 SA-binding pocket are conserved among Arabidopsis NPR1 to NPR4.

**Fig. S8** Yeast two-hybrid interaction between separated LENRV-like and NIMIN1/NIMIN2-binding domain regions of Arabidopsis NPR1 and NPR2, respectively.

**Fig. S9** Chemical structures of salicylic acid and analogs.

**Fig. S10** Effects of salicylic acid analogs on protein-protein interactions of Arabidopsis NPR1, NPR3 and NPR4 in yeast.

**Fig. S11** Effects of salicylic acid analogs on protein-protein interactions of Arabidopsis NPR1, NPR3 and NPR4 in yeast.

**Fig. S12** Formation of hybrid receptors by salicylic acid and structural analogs .

**Fig. S13** Effects of salicylic acid analogs on protein-protein interactions of Arabidopsis NPR3 and NPR4 in yeast.

**Fig. S14** Effects of salicylic acid analogs on *PR-1a* and *NIMIN1* promoter activity in transgenic tobacco plants.

**Table S1** Primers used for construction of full ORF and partial *NPR* cDNA clones.

**Table S2** Primers used for construction of mutant *NPR* cDNA clones.

**Table S3** Primers used for construction of TGA factor encoding cDNA clones.

**Methods S1** Detailed description of methods.

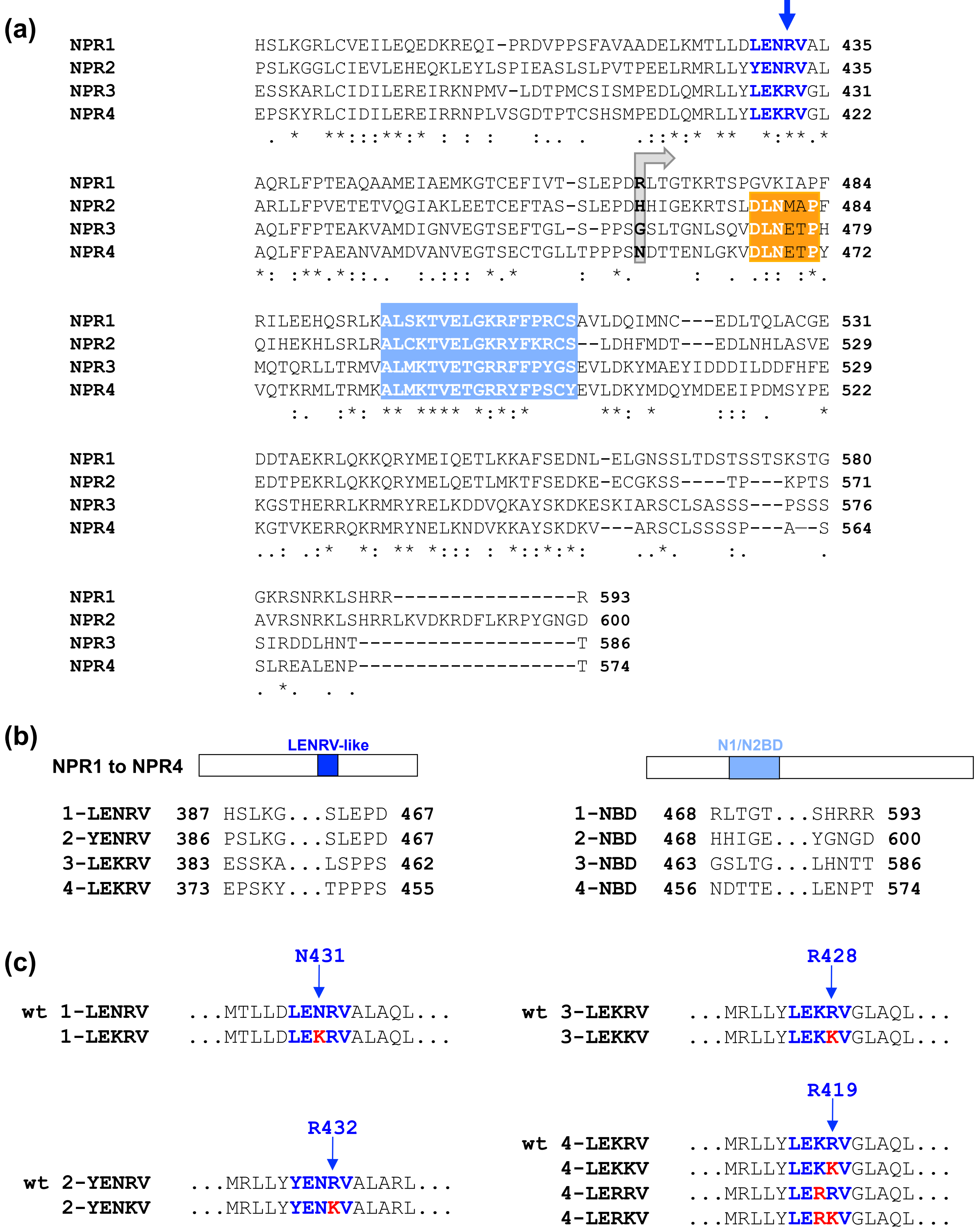


**Fig. S1** Sequence alignment of C-terminal regions of Arabidopsis NPR1 to NPR4 and mutants used in Y2H analyses. (a) Multiple sequence alignment of C-terminal regions of NPR1 to NPR4. Conserved domains, LENRV-like motif with the critical arginine (in navy-blue), DLNxxP-type EAR motif (in orange), and NIMIN1/NIMIN2-binding domain (N1/N2BD; in light blue), are highlighted. C-terminal subclones encompassing the LENRV-like regions and the C-termini of NPR1 to NPR4 start with the first amino acid shown. The grey arrow marks the start of C-terminal subclones spanning the N1/N2BD regions. (b) Extensions of LENRV-like and N1/N2BD subregions of NPR1 to NPR4 as analyzed in Y2H interaction assays. (c) Mutants of NPR1 to NPR4 analyzed in Y2H interactions. Mutated residues are shown together with the respective wild-type (wt) sequence. Mutant subclones have the same extensions as their wild-type correspondents.

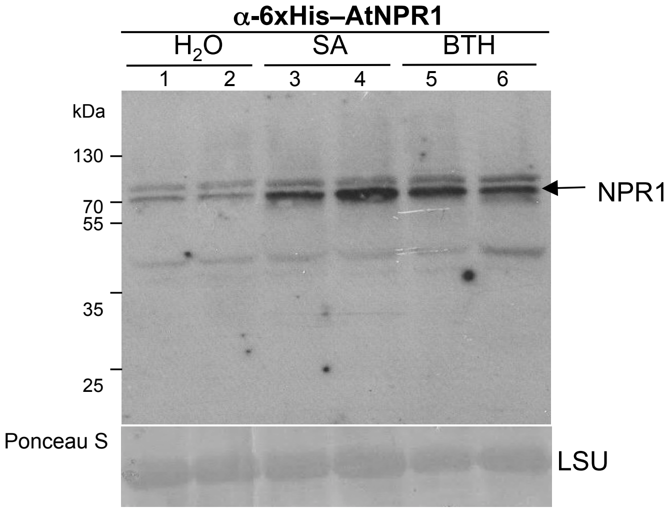


**Fig. S2** Accumulation of Arabidopsis NPR1 in leaf tissue. Arabidopsis Col-0 plants were sprayed with $H_2O$, 1mM SA or 0.3mM BTH. Leaf tissue was harvested from 2 plants for each treatment after 14 hr. Immunodetection was performed with an antiserum directed against 6xHis–AtNPR1 fusion protein. Staining of the large subunit of RUBISCO (LSU) with Ponceau S demonstrates equal loading of the nitrocellulose filter.

(a)

(b)

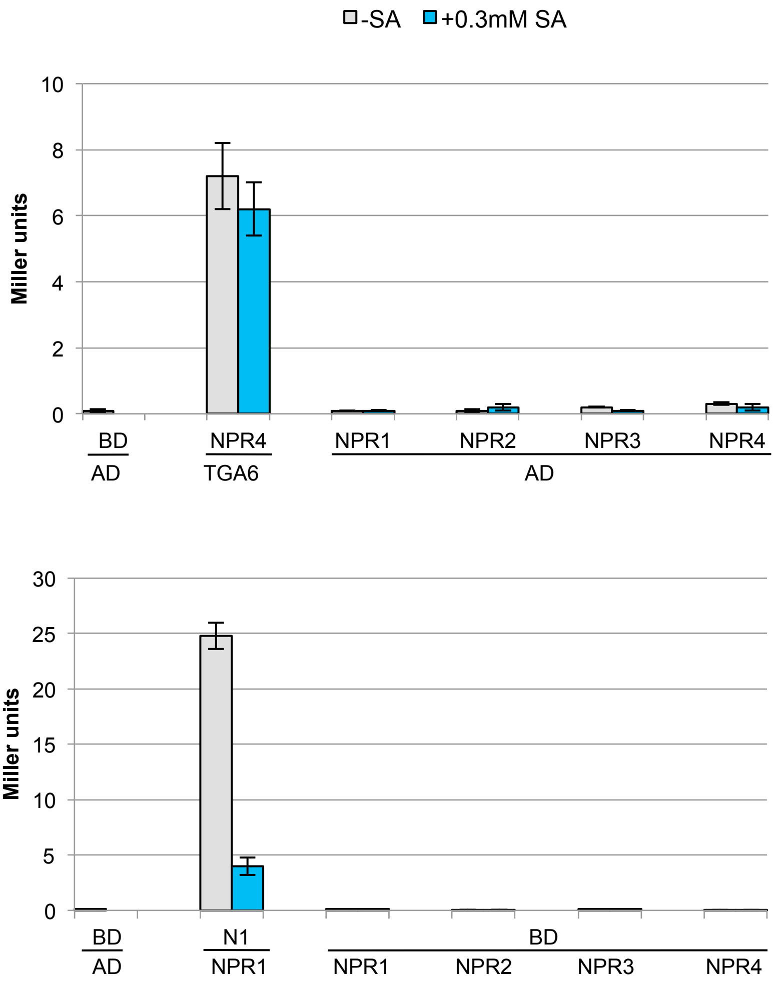


**Fig. S3** Test for autoactivation of Arabidopsis NPR1 to NPR4 in yeast. Transactivation potential of NPR proteins fused to (a) the Gal4 DNA-binding domain (BD) or (b) the Gal4 transactivation domain (AD) was tested in quantitative Y1H assays in absence and presence of salicylic acid (SA). Positive Y2H controls were included to demonstrate activity of BD–NPR4 and AD–NPR1 fusion proteins.

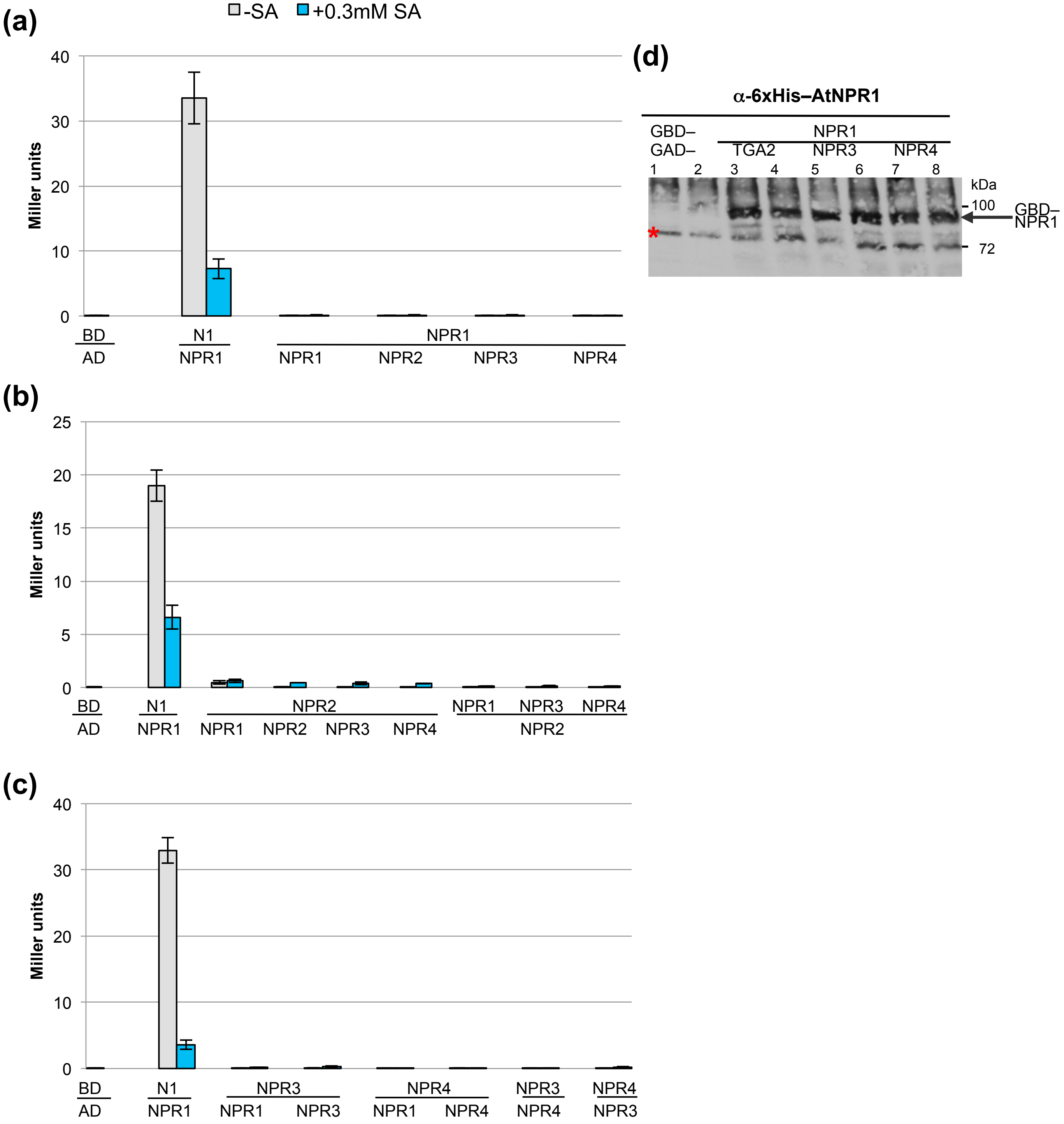


**Fig. S4** Test for homo- and heterodimerization among Arabidopsis NPR1 to NPR4 in yeast. (a,b,c) Dimerization potential among NPR proteins was tested in all combinations with NPR proteins fused to the Gal4 DNA-binding domain (BD) and the Gal4 transactivation domain (AD) in quantitative Y2H assays in absence and presence of salicylic acid (SA). Interaction of BD–NIMIN1 with AD–NPR1 was used as a reference in all test series. (d) Accumulation of GBD–NPR1 in yeast. Protein extracts from 2 independent colonies for each transformation were analyzed by immunodetection with an antiserum directed against 6xHis–AtNPR1. The position of GBD–NPR1 in the gel is marked by an arrow. An unspecific band indicating equal loading of the gel is marked by an asterisk. GBD-NPR1 accumulates in presence of GAD–NPR3 and GAD–NPR4.

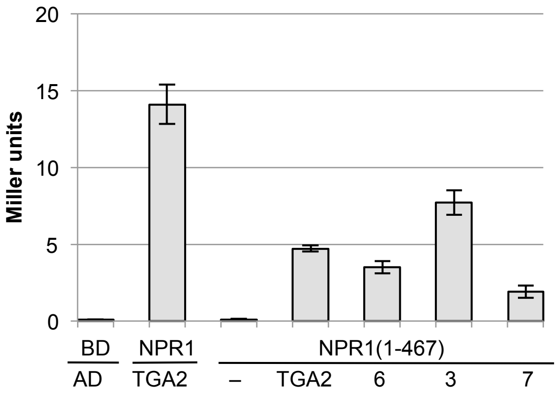


**Fig. S5** Interaction of Arabidopsis TGA transcription factors with truncated Arabidopsis NPR1 in yeast. A C-terminal deletion clone of *NPR1* harboring the BTB/POZ domain and the ankyrin repeats was expressed in fusion with the sequence for the Gal4 DNA-binding domain (BD). Interaction with TGA factors (TGA2, TGA6, TGA3 and TGA7) fused to the Gal4 transactivation domain (AD) was determined in quantitative Y2H assays. Interaction of full-length BD–NPR1 with AD–TGA2 was used as a reference.

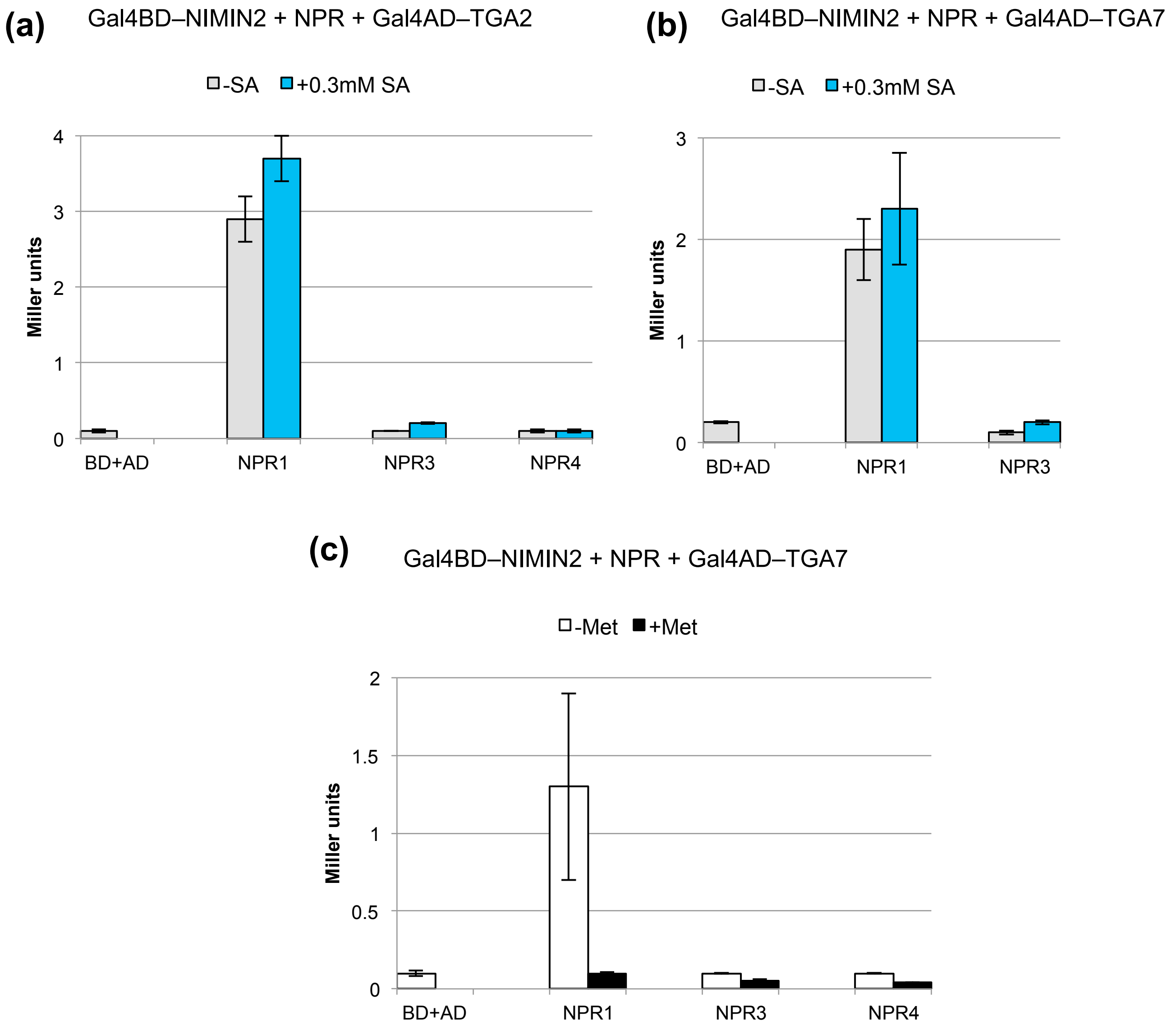


**Fig. S6** Formation of ternary complexes between Arabidopsis NIMIN2, TGA factors and NPR1, NPR3 or NPR4 in yeast. *NIMIN2* was expressed in fusion with the sequence for the Gal4 DNA-binding domain (BD), and *TGAs* were expressed in fusion with the sequence for the Gal4 transactivation domain (AD). *NPRs* were under control of the *MET25* promoter. (a,b) Interaction between Arabidopsis NIMIN2, NPR1, NPR3 or NPR4 and TGA2 or TGA7. Formation of protein complexes was determined in absence of methionine. (c) Interaction between Arabidopsis NIMIN2, NPR1, NPR3 or NPR4 and TGA7. Formation of protein complexes was determined in absence and presence of methionine.

```
                                                                            ↓
                                                                             *
NPR1    HSLKGRLCVEILEQEDKREQI-PRDVPPSFAVAADELKMTLLDLENRVAL 435
NPR2    PSLKGGLCIEVLEHEQKLEYLSPIEASLSLPVTPEELRMRLLYYENRVAL 435
NPR3    ESSKARLCIDILEREIRKNPMV-LDTPMCSISMPEDLQMRLLYLEKRVGL 431
NPR4    EPSKYRLCIDILEREIRRNPLVSGDTPTCSHSMPEDLQMRLLYLEKRVGL 422
         . *  **:::**:* : : :   :.. .    .::*:* **  *:**.*

        *   *  **  *
NPR1    AQRLFPTEAQAAMEIAEMKGTCEFIVT-SLEPDRLTGTKRTSPGVKIAPF 484
NPR2    ARLLFPVETETVQGIAKLEETCEFTAS-SLEPDHHIGEKRTSLDLNMAPF 484
NPR3    AQLFFPTEAKVAMDIGNVEGTSEFTGL-S-PPSGSLTGNLSQVDLNETPH 479
NPR4    AQLFFPAEANVAMDVANVEGTSECTGLLTPPPSNDTTENLGKVDLNETPY 472
        *: :**.*:::.  :.::: *.*     :  *.     :  . .:: :*.

                          ** *   *  ** *
NPR1    RILEEHQSRLKALSKTVELGKRFFPRCSAVLDQIMNC---EDLTQLACGE 531
NPR2    QIHEKHLSRLRALCKTVELGKRYFKRCS--LDHFMDT---EDLNHLASVE 529
NPR3    MQTQRLLTRMVALMKTVETGRRFFPYGSEVLDKYMAEYIDDDILDDFHFE 529
NPR4    VQTKRMLTRMKALMKTVETGRRYFPSCYEVLDKYMDQYMDEEIPDMSYPE 522
           :.  :*: ** ****  *:*:*      **: *     ::: .    *

NPR1    DDTAEKRLQKKQRYMEIQETLKKAFSEDNL-ELGNSSLTDSTSSTSKSTG 580
NPR2    EDTPEKRLQKKQRYMELQETLMKTFSEDKE-ECGKSS----TP---KPTS 571
NPR3    KGSTHERRLKRMRYRELKDDVQKAYSKDKESKIARSCLSASSS---PSSS 576
NPR4    KGTVKERRQKRMRYNELKNDVKKAYSKDKV---ARSCLSSSSP---A--S 564
        ..: .:*  *: ** *::: : *::*:*:    ..*.    :.      .

NPR1    GKRSNRKLSHRR----------------R 593
NPR2    AVRSNRKLSHRRLKVDKRDFLKRPYGNGD 600
NPR3    SIRDDLHNT-------------------T 586
NPR4    SLREALENP-------------------T 574
        . *.  . .
```

**Fig. S7** Amino acid residues forming the NPR4 SA-binding pocket are conserved among Arabidopsis NPR1 to NPR4. Residues forming the SA-binding pocket as determined by X-ray crystallography for SA-bound NPR4 (Wang *et al.,* 2020) are marked by an arrrow (NPR4 R419) and by asterisks (in navy-blue) above the sequence alignment as shown in Fig. S1a. Conserved domains as defined by Maier *et al.* (2011) and Neeley *et al.* (2019) are highlighted (LENRV-like motif with the critical arginine in navy-blue, NIMIN1/ NIMIN2- binding domain (N1/N2BD) in light blue). Red asterisks under the sequence alignment denote amino acids forming the NPR4 SA-binding pocket that are identical among all NPR1 to NPR4.

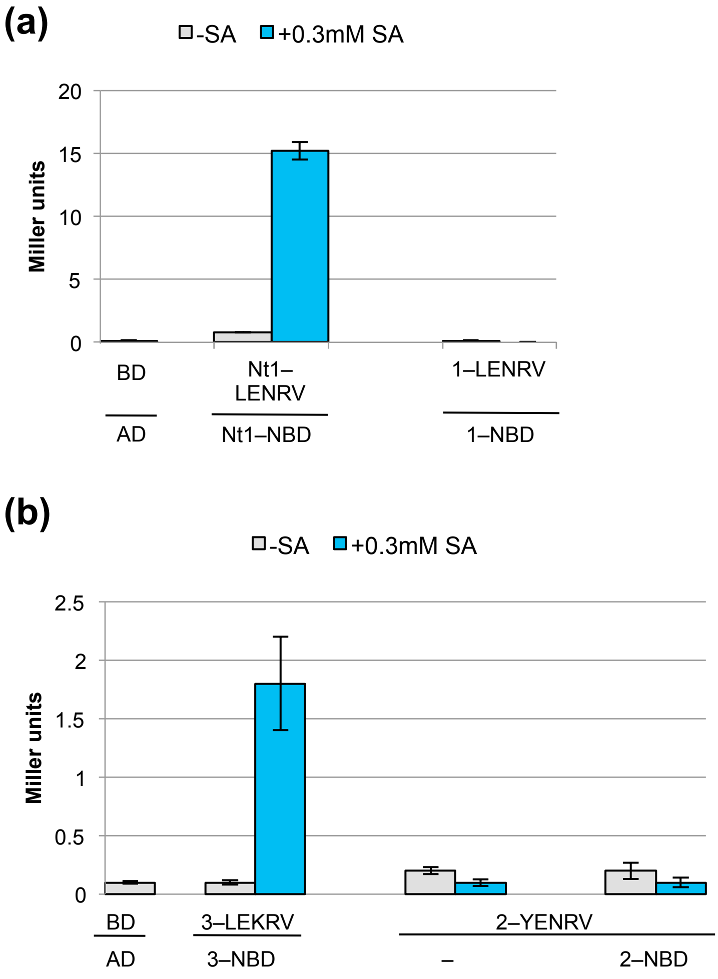


**Fig. S8** Yeast two-hybrid interaction between separated LENRV-like and NIMIN1/NIMIN2-binding domain regions of Arabidopsis NPR1 and NPR2, respectively. Interactions were determined in absence and presence of salicylic acid (SA) in quantitative Y2H assays. (a) Interaction of separated LENRV and N1/N2BD regions of NPR1. Interaction of BD–NtNPR1(Nt1)-LENRV with AD–NtNPR1-N1/N2BD (NtN1–NBD) served as a positive control (Neeley *et al.,* 2019). (b) Interaction of separated YENRV and N1/N2BD regions of NPR2. Interaction of BD–NPR3-LEKRV with AD–NPR3-N1/N2BD served as a positive control. NBD, N1/N2BD region.

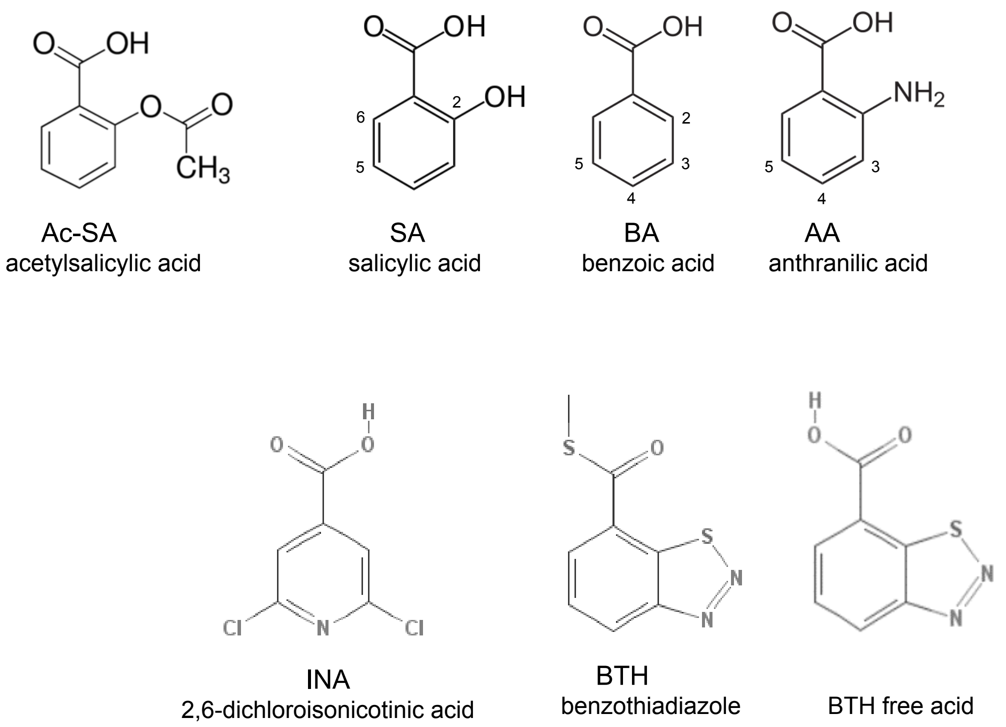


**Fig. S9** Chemical structures of salicylic acid and analogs.

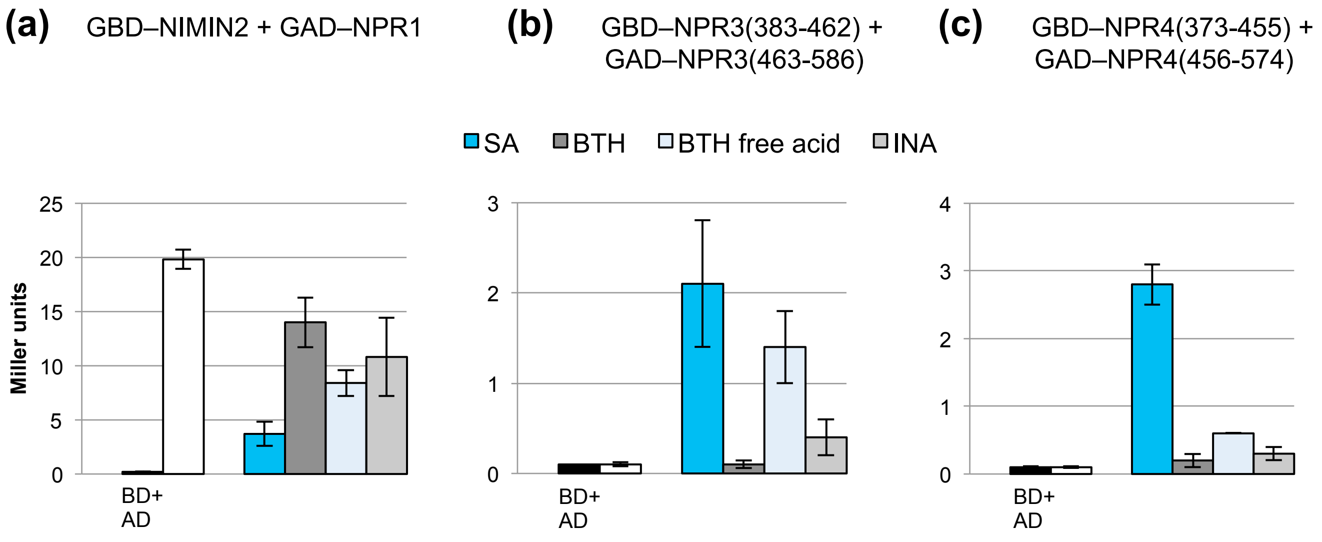


**Fig. S10** Effects of salicylic acid analogs on protein-protein interactions of Arabidopsis NPR1, NPR3 and NPR4 in yeast. Quantitative Y2H assays were conducted with cells grown without addition of chemicals or with cells grown in medium supplemented with SA or analogs. Concentration of chemicals was 300µM. Chemicals were solved in DMSO. The final DMSO concentration in yeast growth medium was 0.5%. (a) Interaction of NIMIN2 with NPR1. (b) Interaction of NPR3(383-462) with NPR3(463-586). (c) Interaction of NPR4(373-455) with NPR4(456-574). SA, salicylic acid; BTH, benzothiadiazole; INA, 2,6-dichloroisonicotinic acid.

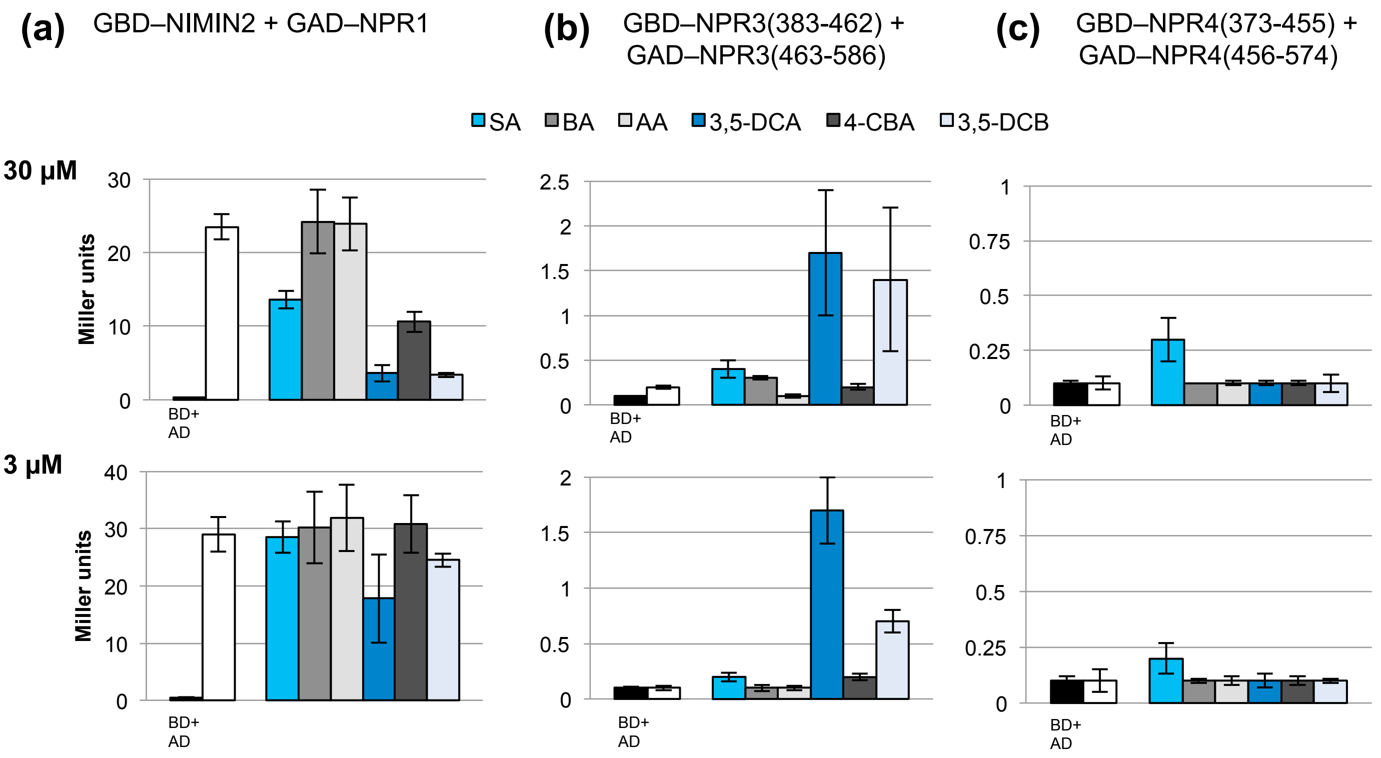


**Fig. S11** Effects of salicylic acid analogs on protein-protein interactions of Arabidopsis NPR1, NPR3 and NPR4 in yeast. Quantitative Y2H assays were conducted with cells grown without addition of chemicals or with cells grown in medium supplemented with SA or analogs. Concentration of chemicals was 30µM or 3µM as indicated. Chemicals were solved in DMSO. The final DMSO concentration in yeast growth medium was 0.5%. (a) Interaction of NIMIN2 with NPR1. (b) Interaction of NPR3(383-462) with NPR3(463-586). (c) Interaction of NPR4(373-455) with NPR4(456-574). SA, salicylic acid; BA, benzoic acid; AA, anthranilic acid; 3,5-DCA, 3,5-dichloroanthranilic acid; 4-CBA, 4-chlorobenzoic acid; 3,5-DCB, 3,5-dichlorobenzoic acid.

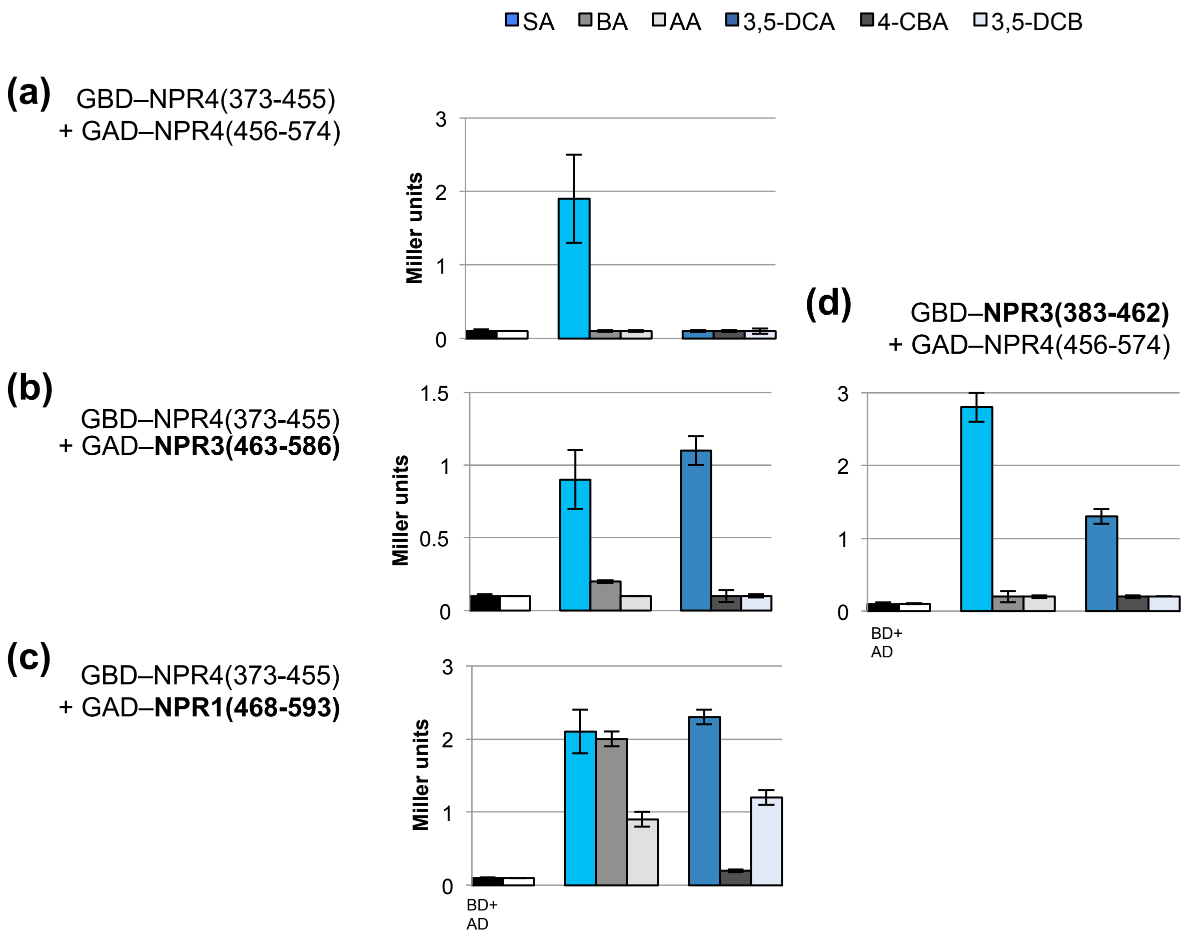


**Fig. S12** Formation of hybrid receptors by salicylic acid and structural analogs. Quantitative Y2H assays were conducted with cells grown without addition of chemicals or with cells grown in medium supplemented with SA or analogs. Concentration of chemicals was 300µM (SA, BA, AA) or 30µM (3,5-DCA , 4-CBA, 3,5-DCB). Chemicals were solved in DMSO. The final DMSO concentration in yeast growth medium was 0.5%. (a-c) The NPR4 LEKRV-region (aa 373-455) was expressed as GBD fusion protein together with GAD–NPR4(456-574) (a), with GAD–NPR3(463-586) (b), and with GAD–NPR1(468-593) (c) in yeast cells. (d) The NPR3 LEKRV-region (aa 383-462) was expressed as GBD fusion protein together with GAD–NPR4(456-574). SA, salicylic acid; BA, benzoic acid; AA, anthranilic acid; 3,5-DCA, 3,5-dichloroanthranilic acid; 4-CBA, 4-chlorobenzoic acid; 3,5-DCB, 3,5-dichlorobenzoic acid.

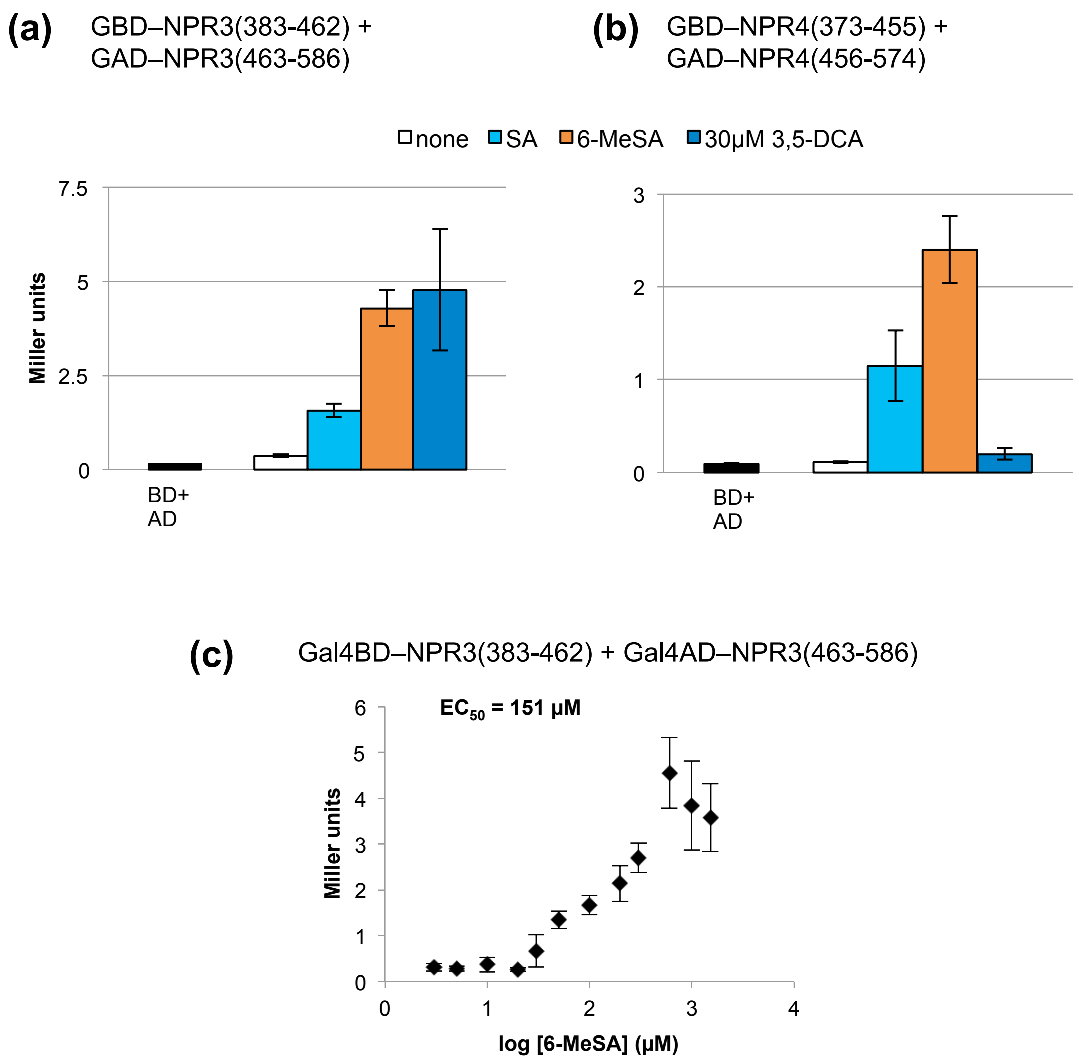


**Fig. S13** Effects of salicylic acid analogs on protein-protein interactions of Arabidopsis NPR3 and NPR4 in yeast. Quantitative Y2H assays were conducted with cells grown without addition of chemicals or with cells grown in medium supplemented with SA or analogs. Concentration of chemicals was 300µM apart from 3,5-DCA (30µM). Chemicals were solved in DMSO. The final DMSO concentration in yeast growth medium was 0.1%. (a) Interaction of NPR3(383-462) with NPR3(463-586). (b) Interaction of NPR4(373-455) with NPR4(456-574). (c) Determination of the half-maximal 6-methylsalicylic acid concentration for NPR3(383-462)-NPR3(463-586) interaction. SA, salicylic acid; 6-MeSA, 6-methylsalicylic acid; 3,5-DCA, 3,5-dichloroanthranilic acid.

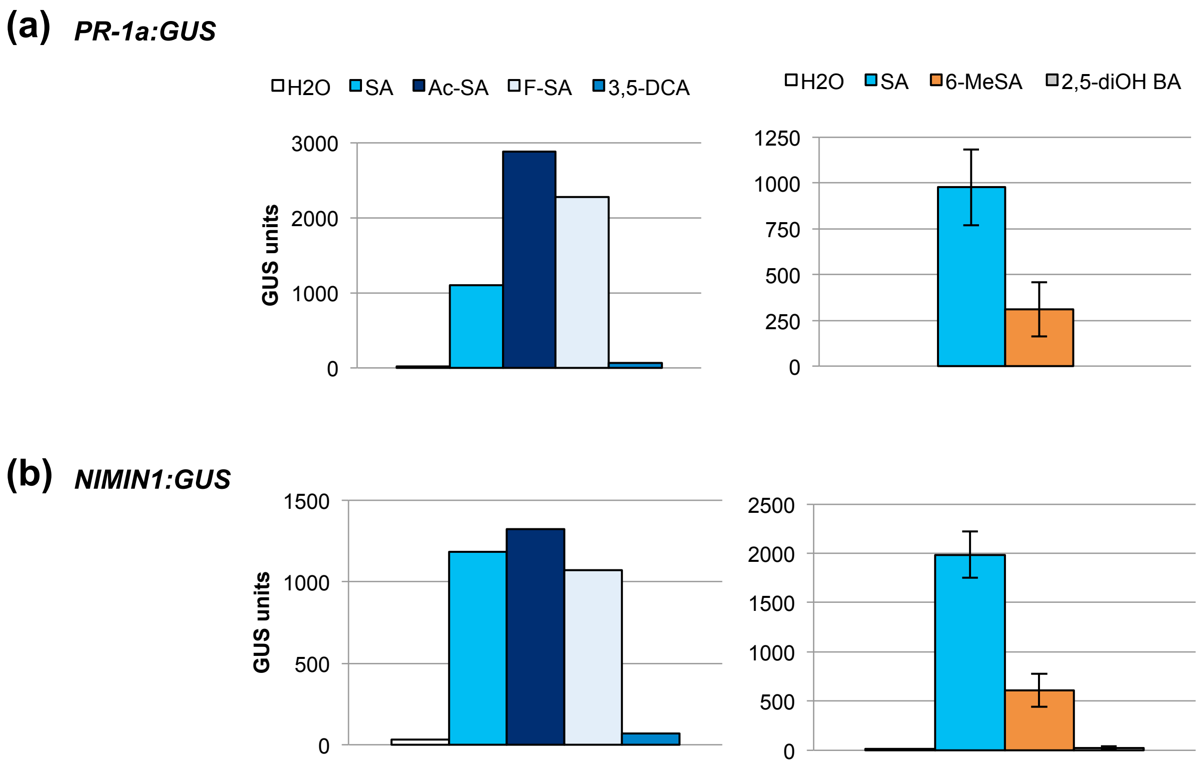


**Fig. S14** Effects of salicylic acid analogs on *PR-1a* and *NIMIN1* promoter activity in transgenic tobacco plants. Leaf discs from transgenic plants harboring *-1533PR1a:GUS* or *NIMIN1:GUS* gene constructs were floated on solutions supplemented with chemicals as indicated (concentration 1mM and 0.1mM for 3,5-DCA; 0.3% DMSO). Average GUS activities from two experiments with different plants or from three experiments with different plants ± standard deviation are shown. (a) Effects on the *PR-1a* pomoter. (b) Effects on the *NIMIN1* promoter. SA, salicylic acid; Ac-SA, acetylsalicylic acid; F-SA, 5-fluorosalicylic acid; 3,5-DCA, 3,5-dichloroanthranilic acid; 6-MeSA, 6-methylsalicylic acid; 2,5-diOH BA, 2,5-dihydroxybenzoic acid.

**Table S1** Primers used for construction of full ORF and partial *NPR* cDNA clones.

| Construct | Primer name | Sequence 5´ to 3´ |
|---|---|---|
| **NPR1**<br>LENRV region | AtNPR1-18<br>AtNPR1-19 | AAGGATCCATATGCATTCTCTCAAAGGCCGAC<br>TTGAGCTCGTCGACTCAGTCAGGCTCGAGGCTAGTC |
| N1/N2BD | AtNPR1-20<br>AtNPR1-21 | AAGGATCCATATGCGTCTCACTGGTACGAAG<br>AAGAGCTCGAGTCAGTCGACCCGACGACGATGAGAGAG |
| **NPR2** exon 1 | AtNPR2-1 | TTGGATCCATATGGCCACCACCACCACCACCACCACCGCTAG |
| 5´-end exon 2 | AtNPR2-13<br>AtNPR2-14 | AAGGATCCATATGTCGCGACAGTTCTTGGAAATTGTAGAC<br>TTGGATCCGGTGGCTCTAGACAGAGCGC |
| exon 2 internal | AtNPR2-15<br>AtNPR2-16 | GCTTTTGGATAGATGCATAG<br>CCTCTATGCATAATCCGCC |
| 3´-end exon 2 | AtNPR2-3<br>AtNPR2-7 | AAGGATCCATGATCTCGTGATGATGGCCACC<br>TCGGTTTTCATAATAGAGCAACCTCATCCTC |
| exon 3 | AtNPR2-8<br><br>AtNPR2-10 | GAGGATGAGGTTGCTCTATTATGAAAACCGAGTTGCACTTGCTCGACTTCTCTTT CCAG<br>GGTTTTACAAAGTGCTCTTAGTCTACTCAAATG |
| exon 4 | AtNPR2-9<br><br>AtNPR2-6 | GCATTTGAGTAGACTAAGAGCACTTTGTAAAACCGTGGAACTGGGGAAACGCT ACTTCAAACG<br>AAGTCGACTTACTCGAGATCCCCGTTCCCGTAAGGTCG |
| YERNV region | AtNPR2-5 | AAGTCGACTTACTCGAGATCAGGCTCGAGACTAGAAGC |
| N1/N2BD | AtNPR2-4 | AAGGATCCAGATGCATCACATTGGTGAAAAGCGG |
| C-terminus | AtNPR2-11<br>AtNPR2-12 | GCGAGTTTACAGCTTCTAGTCTCGAGCCTGATCATCACATTGGTGAAAAGCGG<br>GATCAGGCTCGAGACTAGAAGCTGTAAACTCGC |
| **NPR3**<br>N-terminus | AtNPR3-1<br>AtNPR3-9 | AAGGATCCATATGGCTACTTTGACTGAGCC<br>AAGGATCCGCTACTTCGGGAGGAACTTCC |
| LEKRV region | AtNPR3-3<br>AtNPR3-4 | TTGGATCCATATGGAATCTAGTAAAGCCAGG<br>TTGTCGACTTACTCGAGCGAAGGAGGTGACAAACCCG |
| N1/N2BD | AtNPR3-5<br>AtNPR3-6 | CCGGATCCATATGAGTGGGTTAACCGGAAAC<br>CCGTCGACTTACTCGAGTGTTGTGTTGTGCAGGTC |
| *Bam*HI 3´-end | AtNPR3-10<br>AtNPR3-11 | TTGGATCCATATGAACCCCATGGTTCTAGATACACC<br>CCCTGCAGGTCGACTCAGGATCCTGTTGTGTTGTGCAGGTCATCTC |
| **NPR4**<br>exon 1 | AtNPR4-1<br>AtNPR4-13 | TTGGATCCATATGGCTGCAACTGCAATAGAGCC<br>TACTAGTGACTTCTCAACATAGTTACGAAGCTTCCGCTGAAATGACGAAACAAG ATCCGGG |
| exon 2 | AtNPR4-14<br>AtNPR4-15 | AAGGATCCATATGTCACTAGTAGAGAATGTTCTTCCTATCCTCTTAGTTGCG<br>CCCTTTCCAAGATATCGATGCATAAGC |
| LEKRV region | AtNPR4-5<br>AtNPR4-6 | TTGGATCCATATGGAACCTAGTAAATACCGC<br>AAGTCGACTTACTCGAGTGATGGAGGTGGAGTTAG |
| N1/N2BD | AtNPR4-7<br>AtNPR4-8 | TTGGATCCATATGAATGATACAACTGAAAACTTGGG<br>AAGTCGACTTACTCGAGTGTTGGATTCTCTAAGGC |
| *Bam*HI 3´-end | AtNPR4-3<br>AtNPR4-17 | AAGGATCCATCGCTTATGCATCGATATCTTGG<br>CGGGATCCTGTTGGATTCTCTAAGGC |

**Table S2** Primers used for construction of mutant *NPR* cDNA clones.

| Construct | Primer name | Sequence 5´ to 3´ |
|---|---|---|
| **NPR1** N431K<br>LEKRV | AtNPR1-27<br>AtNPR1-28 | GACGCTGCTCGATCTTGAAAAAAGAGTTGCACTTGCTCAACG<br>CGTTGAGCAAGTGCAACTCTTTTTTCAAGATCGAGCAGCGTC |
| **NPR2** R432K<br>YENKV | AtNPR2-20<br>AtNPR2-22<br>AtNPR2-21<br>AtNPR2-23 | GCGGATGTTAACCTTAGAAATCCG<br>AGAGAAGTCGAGCAAGTGCAACTTTGTTTTCATAATAGAGCAACC<br>GGTTGCTCTATTATGAAAACAAAGTTGCACTTGCTCGACTTCTCT<br>CTACGCTAGCAAGATGATTCAAGTC |
| **NPR3** R428K<br>LEKKV | AtNPR3-21<br>AtNPR3-22 | GAGACTGTTGTACCTAGAAAAGAAAGTGGGGCTTGCTCAG<br>CTGAGCAAGCCCCACTTTCTTTTCTAGGTACAACAGTCTC |
| **NPR4** R419K<br>LEKKV | AtNPR4-9<br>AtNPR4-10 | GAGGTTGTTATACTTAGAAAAGAAAGTGGGACTTGCTCAG<br>CTGAGCAAGTCCCACTTTCTTTTCTAAGTATAACAACCTC |
| K418R<br>LERRV | AtNPR4-11<br>AtNPR4-12 | GAGGTTGTTATACTTAGAAAGGCGAGTGGGACTTGCTCAG<br>CTGAGCAAGTCCCACTCGCCTTTCTAAGTATAACAACCTC |
| K418R R419K<br>LERKV | AtNPR4-15<br>AtNPR4-16 | GAGGTTGTTATACTTAGAAAGGAAAGTGGGACTTGCTCAG<br>CTGAGCAAGTCCCACTTTCCTTTCTAAGTATAACAACCTC |

**Table S3** Primers used for construction of TGA factor encoding cDNA clones.

| Construct | Primer name | Sequence 5´ to 3´ |
|---|---|---|
| **TGA3** | AtTGA3-1<br>AtTGA3-2 | CCGGATCCATATGGAGATGATGAGCTCTTC<br>AAGGATCCAGTGTGTTCTCGTGGACGAGC |
| **TGA7** | AtTGA7-1<br>AtTGA7-2 | AAGGATCCATATGATGAGTTCTTCTTCTCCAAC<br>AAGGATCCAGTTGGTTCTTGTGGACGAGC |

**Methods S1** Detailed description of methods.

**Construction of full-length and partial *NPR1*, *NPR2*, *NPR3* and *NPR4* cDNA clones**

Subregions of *NPR* genes used in Y2H are depicted in Fig. S1a and b, and primers used in PCR amplifications are listed in Table S1.

Clones encoding the LENRV and N1/N2BD regions of NPR1 and analogous partial cDNA clones for *NPR2* to *NPR4* were constructed following the strategy employed previously for tobacco *NPR1* subclones (Neeley *et al.*, 2019). In short, the C-terminal third of Arabidopsis NPR1 (aa 387-593; Stos-Zweifel *et al.*, 2018) was split in the middle between the LENRV motif and the N1/N2BD resulting in subregions of amino acids 387 to 467 (LENRV region) and 468 to 593 (N1/N2BD region). The PCR-amplified sequences were cloned as *Bam*HI/*Sal*I fragment in pGBT9 (LENRV region) and as *Bam*HI/*Xho*I fragment in *Bam*HI/*Sal*I cleaved pGAD424 (N1/N2BD).

A full ORF cDNA clone for *NPR2* was obtained by PCR from genomic Arabidopsis Col-0 DNA. First, a cDNA clone comprising the C-terminal third from amino acids 386 to 600 was constructed. Three PCR products were generated spanning the C-terminal end of exon 2, whole exon 3 with a 5′-overhang of exon 2 sequence, and whole exon 4 with a 5′-overhang of exon 3 sequence. Exon 2 and 3 fragments, and exon 3 and 4 fragments, respectively, were combined by overlap extension PCR as described by Ho *et al.* (1989). From these clones, the sequences coding for the YENRV and N1/N2BD regions were amplified by PCR and ligated as *Bam*HI/*Sal*I fragments to pGBT9 and pGAD424. The resulting YENRV and N1/N2BD clones were then used as templates in overlap extension PCR performed with complementary primers AtNPR2-11 and AtNPR2-12 spanning the YENRV/N1/N2BD boundary to assemble a cDNA clone encoding the NPR2 C-terminal third. The sequence was cloned as *Bam*HI/*Sal*I fragment in pGBT9, pGAD424 and pUC19. To the pUC19/NPR2(386/600) clone, PCR-amplified sequences encoding the 5′-end of exon 2 and an internal exon 2 fragment were added as *Bam*HI and *Nsi*I fragments, respectively, to yield pUC19/NPR2(187-600). The *NPR2* fragment was cloned in *Bam*HI/*Sal*I cut pGAD424, and the first exon sequence, from the AUG start codon to a unique *Xba*I restriction site in exon 2, was amplified by PCR and added as *Bam*HI/*Xba*I fragment to yield the full ORF *NPR2* cDNA. The *NPR2* cDNA was ligated as *Bam*HI/*Sal*I fragment to pGBT9.

For generating *NPR3* and *NPR4* clones, a full-length *NPR3* cDNA clone (pda09552-RAFL07-17-D18; Riken BRC Experimental Plant Division, Japan; Seki *et al.*, 1998, 2002 ) and a partial *NPR4* cDNA clone (CATMA4a20900/CD269995; Arabidopsis Biological Resource Center, The Ohio State University; Hilson *et al.*, 2004) were used as PCR templates to obtain the sequences coding for truncated proteins encompassing the LEKRV, N1/N2BD and C-terminal regions. The sequences were cloned as *Bam*HI/*Sal*I fragments in vectors pGBT9 and pGAD424. To obtain a full ORF *NPR3* cDNA clone, a 0.7 kb fragment encoding the N-terminus was amplified with primers AtNPR3-1 and AtNPR3-9 and ligated as *Bam*HI fragment to pGBT9/NPR3(383-586). To this clone, an internal 0.6 kb *Bgl*II fragment isolated from the RIKEN clone was added. The cDNA was ligated as *Bam*HI/*Pst*I fragment to pGAD424. To generate a cDNA with *Bam*HI ends, the 3′-end of the *NPR3* cDNA was replaced with a PCR-generated *Nco*I/*Pst*I fragment containing a *Bam*HI restriction site 5′ to *Pst*I. To construct a full ORF *NPR4* cDNA, sequences encoding exons 1 and 2 of the *NPR4* gene were amplified from genomic DNA. The exon 1 sequence contained an overhang of exon 2 sequence including a unique *Spe*I restriction site, and the exon 2 sequence crossed a unique *Eco*RV restriction site which is located in the cDNA sequence 5′ to the LEKRV motif. The C-terminal *NPR4* sequence was added to the exon 2 clone as *Eco*RV/*Sal*I fragment, and the combined cDNA was ligated as *Bam*HI/*Sal*I fragment to pGBT9 yielding pGBT9/NPR4(183-574). To this clone, the exon 1 sequence was added as *Bam*HI/*Spe*I fragment. The cDNA was cloned as *Bam*HI/*Sal*I fragment in pGAD424. To generate a cDNA with *Bam*HI ends, the 3′-end was amplified with primers AtNPR4-3 and AtNPR4-17, and the resulting *Bam*HI fragment was ligated to pUC19. To this clone, the 5′-half of *NPR4* was added as *Sma*I/*Eco*RV fragment excised from pGAD424/NPR4. The full ORF cDNA was re-inserted as *Bam*HI fragment in pGAD424 .

For expression of *NPR3* and *NPR4* in Y3H analyses, gene sequences were excised from pGAD424 constructs as *Bam*HI fragments and inserted in the *Bgl*II restriction enzyme site of a modified pBridge vector, designated pBD–NIMIN2/-, under control of the *MET25* promoter (Weigel *et al.*, 2001).

**Construction of mutant *NPR* cDNA clones**

Mutants used in Y2H analyses are depicted in Fig. S1c. Mutations were introduced in *NPR* cDNAs by overlap extension PCR. Primers used in PCR amplifications are listed in Table S2. Mutation NPR2 R432K was introduced in a *Hpa*I/*Nhe*I fragment, and the mutated fragment was integrated in the wt full ORF cDNA clone. For *NPR1*, *NPR3* and *NPR4*, mutations were introduced in the fragments encoding the proteins′ C-terminal thirds. From the resulting clones, the mutant LENRV-like regions were amplified by PCR and ligated to pGBT9.

**Construction of TGA factor encoding cDNA clones**

For cloning *TGA3* and *TGA7*, ORF cDNA clones were obtained from the Arabidopsis Biological Resource Center, The Ohio State University (pUNI51/TGA3, U82565; pUNI51/TGA7, U14616; Yamada *et al.*, 2003). The clones were used as PCR templates to obtain full ORF cDNAs with *Bam*HI ends which were ligated to pGAD424. Primers used in the PCR amplifications are listed in Table S3.

**Accession numbers**

Sequence data from this article can be found in the EMBL/GenBank databases under the following accession numbers:

AF480488 (*NtNPR1*), At1g64280 (*NPR1*), At4g26120 (*NPR2*), At5g45110 (*NPR3*), At4g19660 (*NPR4*), At5g06600 (*TGA2),* At1g22070 (*TGA3*), At3g12250 (*TGA6*), At1g77920 (*TGA7*), At1g02450 (*NIMIN1*), At3g25882 (*NIMIN2*).

**Hilson P, Allemeersch J, Altmann T, Aubourg S, Avon A, Beynon J, Bhalerao RP, Bitton F, Caboche M, Cannoot B *et al.* 2004.** Versatile gene-specific sequence tags for *Arabidopsis* functional genomics: transcript profiling and reverse genetics applications. *Genome Research* **14**: 2176-2189.

**Ho SN, Hunt HD, Horton RM, Pullen JK, Pease LR. 1989.** Site-directed mutagenesis by overlap extension using the polymerase chain reaction. *Gene* **77**: 51-59.

**Maier F, Zwicker S, Hückelhoven A, Meissner M, Funk J, Pfitzner AJP, Pfitzner UM. 2011.** NONEXPRESSOR OF PATHOGENESIS-RELATED PROTEINS1 (NPR1) and some NPR1-related proteins are sensitive to salicylic acid. *Molecular Plant Pathology* **12**: 73-91.

**Neeley D, Konopka E, Straub A, Maier F, Pfitzner AJP, Pfitzner UM. 2019.** Salicylic acid-driven association of LENRV and NIMIN1/NIMIN2 binding domain regions in the C-terminus of tobacco NPR1 transduces SAR signal. *Bioarchives*. doi: 10.1101/543645

**Seki M, Carninci P, Nishiyama Y, Hayashizaki Y, Shinozaki K. 1998.** High-efficiency cloning of Arabidopsis full-length cDNA by biotinylated CAP trapper. *Plant Journal* **15**: 707-720.

**Seki M, Narusaka M, Kamiya A, Ishida J, Satou M, Sakurai T, Nakajima M, Enju A, Akiyama K, Oono Y *et al.* 2002.** Functional annotation of a full-length Arabidopsis cDNA collection. *Science* **296**: 141-145.

**Stos-Zweifel V, Neeley D, Konopka E, Meissner M, Hermann M, Maier F, Haefner V, Pfitzner AJP, Pfitzner UM. 2018.** Tobacco TGA7 mediates gene expression dependent and independent of salicylic acid. *Bioarchives.* doi: 10.1101/341834

**Wang W, Withers J, Li H, Zwack PJ, Rusnac D-M, Shi H, Liu L, Yan S, Hinds TR, Guttman M *et al.* 2020.** Structural basis of salicylic acid perception by *Arabidopsis* NPR proteins. *Nature* **586**: 311-316.

**Weigel RR, Bäuscher C, Pfitzner AJP, Pfitzner UM. 2001.** NIMIN-1, NIMIN-2 and NIMIN-3, members of a novel family of proteins from Arabidopsis that interact with NPR1/NIM1, a key regulator of systemic acquired resistance in plants. *Plant Molecular Biology* **46**: 143-160.

**Yamada K, Lim J, Dale JM, Chen H, Shinn P, Palm CJ, Southwick AM, Wu HC, Kim C, Nguyen M *et al.* 2003.** Empirical analysis of transcriptional activity in the *Arabidopsis* genome. *Science* **302**: 842-846.
